# Functional MRI of large scale activity in behaving mice

**DOI:** 10.1101/2020.04.16.044941

**Authors:** Madalena S. Fonseca, Mattia G. Bergomi, Zachary F. Mainen, Noam Shemesh

**Author notes:** Correspondence to (M. S. F.) or (N. S.). These authors contributed equally (Zachary F. Mainen, Noam Shemesh).

## Abstract

Behaviour involves complex dynamic interactions across many brain regions. Detecting whole-brain activity in mice performing sophisticated behavioural tasks could facilitate insights into distributed processing underlying behaviour, guide local targeting, and help bridge the disparate spatial scales between rodent and human studies. Here, we present a comprehensive approach for recording brain-wide activity with functional magnetic resonance imaging (fMRI) compatible with a wide range of behavioural paradigms and neuroscience questions. We introduce hardware and procedural advances to allow multi-sensory, multi-action behavioural paradigms in the scanner. We identify signal artefacts arising from task-related body movements and propose novel strategies to reduce them. We validate and explore our approach in a 4-odour classical conditioning and a visually-guided operant task, illustrating how it can be used to extract information so far inaccessible to rodent behaviour studies. Our work paves the way for future studies combining fMRI and local circuit techniques during complex behaviour to tackle multi-scale behavioural neuroscience questions.

## INTRODUCTION

Adaptive behaviour requires dynamic interactions between millions of neurons both within local neural circuits and across many brain regions. By offering unprecedented access to neuronal circuits, rodent behavioural studies have provided fundamental insight into how local neuronal activity relates to behaviour. However, a comprehensive understanding of the neural basis of behaviour will depend not only on studying a subset of areas individually (e.g. basal ganglia and motor cortex), but also on observing how interactions between these and other areas give rise to behaviour (e.g. interplay between motor cortex, basal ganglia, cerebellum and thalamus in motor learning and execution).

A brain-wide view would not only provide important insight into behavioural function, but also guide the selection of brain regions to be studied at the level of microcircuits. Invasive studies have traditionally selected targets based on previous literature, leading to the repeated study of the same areas, while overlooking other regions. This would be equivalent to an explorer starting to investigate a new territory by following the existing local cues. But what if the explorer could climb to a high viewpoint, and get a less detailed but more global view of the landscape? Then based on this overview, she could select, in an unbiased way, which places are worth probing. Thus, beyond informing systems-level behavioural theories, a whole-brain approach would offer an unbiased way to select brain regions to be targeted.

By contrast, human cognitive studies routinely access global brain activity but lack detailed descriptions of local processing. This, along with the fact that rodent and human studies typically use different techniques and measure different types of signals, has created a disconnect between the mechanistic findings in rodents and the macroscale findings in humans.

Recent advances in electrophysiology and calcium imaging have extended their spatial scale (Lu et al., 2020; Musall et al., 2019; Sofroniew et al., 2016; Steinmetz et al., 2019; Wekselblatt et al., 2016). However, they are still far from being whole-brain and multi-area simultaneous access is still limited. Furthermore, they are not extendable to humans. Functional ultrasound has emerged as a promising tool combining wide brain coverage with high spatiotemporal resolution (Deffieux et al., 2018; Macé et al., 2018; Rabut et al., 2019), however, acquisitions are still serial in awake rodents (Macé et al., 2018; Sieu et al., 2015; Urban et al., 2014) and invasive in humans (Deffieux et al., 2018). Thus, functional magnetic resonance imaging (fMRI) remains the gold standard for brain-wide simultaneous imaging, allowing activity to be followed non-invasively over time and across species, from mouse to human.

The ability to perform fMRI in awake behaving rodents thus offers both a brain-wide perspective to rodent behavioural neuroscience research and a direct comparison to the signals measured in similar settings in humans. However, despite its enormous potential, rodent behaviour fMRI is still in its infancy (Han et al., 2019; Sakurai et al., 2020; Tabuchi et al., 2002) and the available methods are still far from being comparable to the sophisticated behavioural paradigms and data analysis used in typical non-fMRI rodent and human fMRI studies.

In creating a general method that will be widely useful to the rodent behavioural community and thus work as a bridge to human studies, several challenges remain. First, the method must offer the hardware capabilities and flexibility to replicate the kinds of sophisticated tasks that are commonly used in non-MRI rodent studies to isolate sensory, cognitive and motor processing. Previous fMRI studies in behaving rodents have presented significant hardware developments but remain very low-dimensional (1-2 unimodal sensory cues and 1 motor readout, (Han et al., 2019; Sakurai et al., 2020; Tabuchi et al., 2002). This has resulted in limited ability to interpret the underlying neural activity (e.g. distinguishing sensory processing from reward association or motor execution).

Second, the method must be able to measure neural activity while animals are engaged in motor output. This has remained perhaps the biggest challenge as body and jaw movements, even without changes in head position, are known to cause large artefacts in fMRI signals (Keliris et al., 2007; Tabuchi et al., 2002; Van de Moortele et al., 2002). Given that jaw and other movements (e.g. licking) are highly correlated with task performance, it is crucial that these are corrected especially to avoid false positives. Of the three studies that performed fMRI during active behaviour in rodents, two have only addressed artefacts caused by head movements (Han et al., 2019; Sakurai et al., 2020). The third (Tabuchi et al., 2002) has used an external reference sample, positioned near the head of the animal, to measure and correct global changes in brain intensity caused by jaw and tongue movements. However, this strategy takes into account only global changes, when it is likely that regional effects also occur. Furthermore, it relies on using a big enough external sample positioned within the imaging field of view which is difficult in practice, especially when using more complex behavioural setups. Therefore, existing methods are neither complete nor generalizable in correcting artefactual signals.

Third, the method should include the development of behavioural protocols and task designs that take into account the slow dynamics of the BOLD signal to allow parsing activity related to different behavioural task epochs. This has remained unexplored given the short trial design previously used (2s - Han et al., 2019; Sakurai et al., 2020).

Finally, it must be able to exploit sophisticated neuro-behavioural analysis commonly found in single area rodent studies (e.g. behavioural decoding) and human fMRI studies (e.g. inter-area coupling) to illustrate the breadth of new information that can be extracted from this approach.

Here, we present a comprehensive approach that tackles these challenges enabling the use of behaving rodent fMRI in a broad range of behavioural paradigms to tackle previously inaccessible systems neuroscience questions. Briefly, we begin by describing an MRI-compatible behaviour setup with multiple sensory cues and motor readouts, and present protocols to train mice to perform sophisticated reward-guided tasks in the scanner. We identify and investigate signal artefacts that correlate with body movements and present novel strategies to reduce them. We start by validating our approach and correction strategies by showing sensory, motor, and reward correlates of mice performing a 4 odour-guided classical conditioning task. Our results are consistent with previous literature on classical conditioning, while highlighting a number of additional areas that have either remained unexplored or only studied in other contexts. Finally, we demonstrate the generality of our approach in an operant conditioning task with higher motor complexity and task flexibility. We further illustrate how a combination of analysis strategies can be used to extract information that has so far been inaccessible to rodent behavioural studies: brain-wide maps of task-related activity that can be used to guide more local investigations, comparisons of task event discriminability across multiple cortical and subcortical areas, and information on inter-region functional coupling.

## RESULTS

### An MRI compatible setup with multiple sensory inputs and behavioural outputs

The strong magnetic field and the restricted space in the scanner means that hardware solutions typically used for rodent head-fixation (i.e. metal head plates) and behavioural quantification (i.e. large metallic components) are not compatible with MR imaging. MRI compatible behavioural setups exist (Han et al., 2019; Sakurai et al., 2020) but have so far been limited to a single readout measurement (one port licking) and maximum 2 sensory stimuli. We aimed to develop a setup that, by allowing multiple sensory stimuli and behavioural readout measures, enabled higher complexity and was thus compatible with a broader range of tasks and neuroscientific questions. Combining fibre optics, pressure sensing, and 3D printed plastic parts, we developed an MRI compatible setup (**Figure 1**) that includes a head-fixation system, multiple sensory delivery systems for olfactory (up to 8 different odours) and visual stimuli (at least 2) and three separate behavioural measures (licking, right lever press, left lever press).

**Figure 1.**
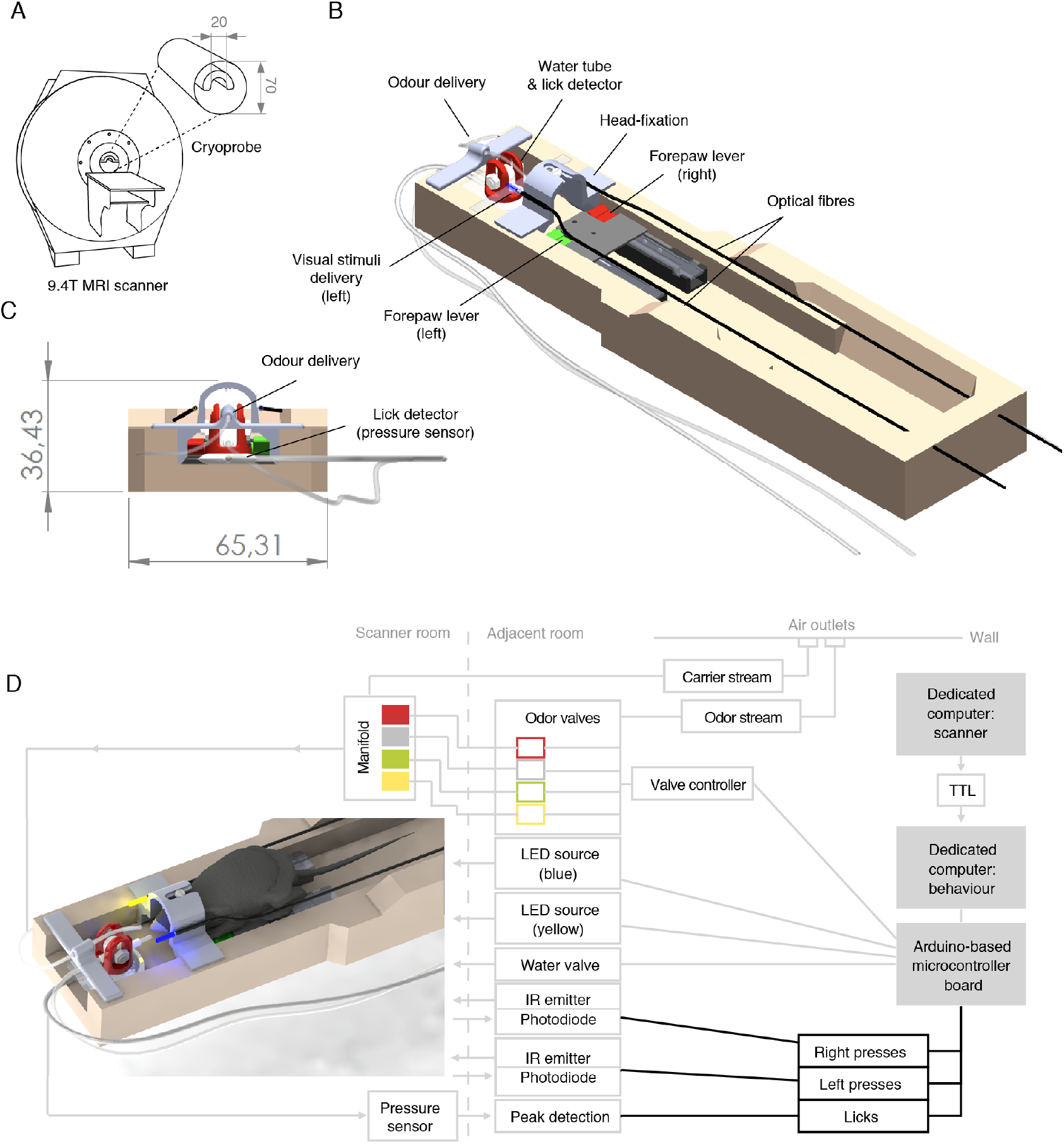
MRI-compatible behaviour setup for head-fixed mice. **(A)** Schematic of the scanner and coil used for MRI imaging. **(B)** Schematic of the behavioural setup, including a head-fixation system, a water delivery tube, a detector for licking behaviour, a detector for right (red) and left (green) forepaw lever presses, an odour delivery tube and two optic fibres delivering blue (left) and yellow (right) light stimuli. **(C)** Front view. **(D)** Upper view showing the mouse position (left) and wiring diagram (right) of the hardware involved. All the electronic and metal components were positioned at a safe distance from the scanner and connected to the setup through long-range connections. Lever pressing was detected using two infra-red beam break systems delivered and sensed via long-range optic fibres. Licking was detected using a pressure sensor that detected lick tube vibration via an air pad positioned under the tube holder. Olfactory stimuli were delivered through a custom built olfactometer connected to a manifold in the scanner that quickly routed the mixed air stream to the animal’s nose via a teflon tube. Visual stimuli were delivered via optic fibres connected to fibre-coupled LED sources. Water was delivered using a distant water valve. All systems connected to an Arduino-based board that was controlled by a dedicated computer using Arduino software, Python and Bonsai. Dimensions are shown in mm.

### Four-odour classical conditioning task in the scanner

Using this setup (**Figure 2A**), we first trained mice in a classical conditioning task (**Figure 2B**), where mice learned to associate different odours (conditioned stimuli, CS) with biologically relevant outcomes (unconditioned stimuli, US), such as the presence or absence of water rewards. We chose classical conditioning as it is well studied and fast to train (J. Y. Cohen et al., 2012, 2015; Matias et al., 2017; Tian & Uchida, 2015). This made it ideally suited for our purpose of optimising and validating our training and imaging methods. We used four different odours, two of which were paired with water reward (CS+), and two of which were not rewarded (CS-) (**Figure 2B**). This was done so that we could later discriminate brain activity related to both odour identity and reward association.

**Figure 2.**
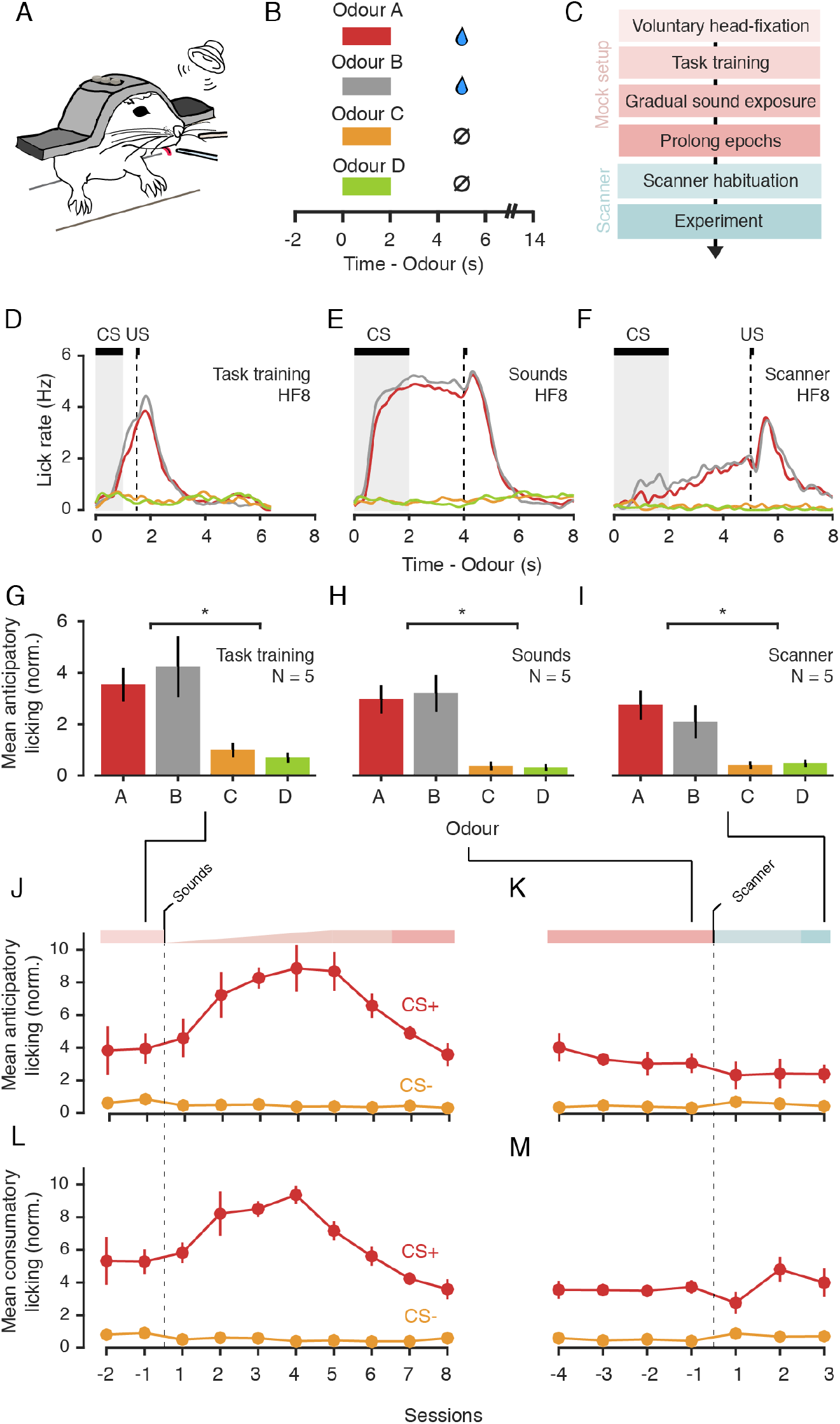
Four-odour classical conditioning task and acclimation procedure. **(A)** Behavioural setup. **(B)** Task structure. A trial started with the delivery of one out of four possible odours (Conditioned Stimuli, CS), randomly selected. Two of the odours (A and B for group 1, N = 3, or C and D for group 2, N = 2, termed CS+) were followed deterministically by a water reward (US) delivered after a fixed delay (trace period) of 3s from odour offset. For the other two odours (CS-), reward was not delivered, but the trial structure was the same. After the outcome there was a variable inter-trial interval (6-9s), after which a new trial began. **(C)** Training procedure. Mice were first trained to voluntarily enter the head-fixation apparatus. Next, mice underwent task training involving repeated CS-US pairings. Once mice learned the task (see D), a mixture of scanner sounds were gradually introduced over 5 sessions and maintained at maximum loudness for at least another 7 sessions. Mice then transitioned to the scanner where they were exposed to the real scanner sounds and its vibration before the experiment. **(D-F)** Averaged lick trace split by odour for a representative mouse (HF8) for the session before sound introduction (D), before transitioning to the scanner (E) or with functional imaging (F). **(G-I)** Mean anticipatory licking (750 ms window before outcome), averaged across mice (N = 5), for the last session of task training (G), sound exposure (H) or scanner exposure (I). **(J-K)** Mean anticipatory lick rate, averaged across mice (N = 5) for CS+ (red) and CS- (grey) trials over sessions aligned on sound exposure (J) or aligned on transition to the scanner (K). **(L-M)** Same as J-K but for the consummatory lick rate (100 to 850 ms from outcome). Error bars represent SEM across mice. To ensure comparability across sessions and across different lick detectors (see Methods), the mean lick rates shown in G-M were normalized to the overall lick rate recorded in the entire session. *p<0.05, paired t-test.

Mice learned the stimulus-outcome association, evidenced by the significantly higher licking rates in the delay period after odours that were paired with rewards relative to odours that were not (normalized lick rate: CS+: 3.94 ± 2.08, CS-: 0.85 ± 0.52, mean ± SD, t(4) = 4.14, p < 0.05, N = 5 mice, **Figures 2D** and **2G**). Importantly, licking began before outcome delivery (anticipatory licking) showing that mice learned the predictive meaning of each odour (**Figure 2D**).

A major challenge in performing fMRI in a behaving mouse is that the scanner is an intrinsically stressful environment. This is due to its restricted space, loud (∼120dB) and variable noises, and strong vibrations. Besides being a concern for animal welfare, excessive stress can impair learning and task performance (de Quervain et al., 1998; Diamond et al., 1994; Graham et al., 2010; Hölscher, 1999; Kaneto, 1997; Kim et al., 2001) and lead to poor image quality due to head motion (Harris et al., 2015; King et al., 2005).

Exposing animals to MRI sounds ahead of imaging has been shown to be effective at decreasing levels of stress in awake non-behaving rodents (Desai et al., 2011; Ferenczi et al., 2016; Harris et al., 2015; King et al., 2005), with longer protocols being the most effective (Harris et al., 2015). Existing protocols for awake behaving mice have used this approach but for a short period (2-6 days) (Han et al., 2019; Sakurai et al., 2020), raising the possibility that, even if lowerer, stress is not being minimized. This may be compatible with the performance of simple tasks, but it is likely problematic for more complex designs given that stress has been shown to render behaviour insensitive to changes in outcome value and resistant to changes in task contingencies (Dias-Ferreira et al., 2009; Graham et al., 2010).

In an effort to create a method that was both compatible with the sophisticated tasks used by the non-MRI rodent community and maximized their comparability, we chose a longer (30-40 days) and more gradual procedure (**Figure 2C,** see **Methods**) that aimed to minimize stress at every stage of training. This protocol differed from previous methods (Han et al., 2019; Sakurai et al., 2020) in four main aspects. First, to minimize any restraint-related stress, we trained mice to voluntarily enter the head-fixation apparatus (8-10 days). This avoided the stress caused by the experimenter restraining and manually positioning the mouse and ensured that mice were fully comfortable before entering the next stage of training. Second, mice were exposed to MRI sounds recorded from the scanner very gradually (increasing in volume over 5 days), only once they were proficient in the task, and before increasing task epochs (**Figure 2C**). This ensured that mice were exposed to only small amounts of stress each day and during a period where they were maximally engaged in the task (at maximum reward rate). Third, we maintained sounds at the highest level over at least 7 more days, in a total of at least 12 days of sound exposure, following previous reports that longer periods of exposure are more effective at minimizing stress (Harris et al., 2015). Fourth, we habituated mice not only to the scanner sounds, but also the vibrations produced in the scanner. Given that these vibrations are hard to replicate in a mock setup, we also trained mice in the scanner before starting experiments.

We expected behaviour to be affected by sounds but normalize over the course of continual exposure. Typically, on the first day of sound exposure, mice interrupted licking at sound onset and after a few seconds resumed licking (data not shown). Licking rate to rewarded odours increased as sound levels increased, both in the anticipatory phase (**Figure 2J**, days 1-5) and after outcome delivery (**Figure L,** days 1-5). It took 7-8 days of repeated exposure for licking to recover to baseline levels (**Figure 2J** and **2L)**.

Once sounds were at the highest level, task epochs were gradually increased to reach the final configuration (odour presentation for 2s, delay from odour offset to outcome (trace) of 3s, and a variable inter-trial interval of 6-9s). This ensured that task epochs were as separable as possible and thus were distinguishable in our fMRI measurements. Increasing the odour and trace periods led initially to mice consistently licking from odour presentation to outcome (**Figures 2E**). Over training, the overall rate of anticipatory licking decreased and started progressively later in the trace period (**Figures 2F** and **2I**), consistent with mice learning to anticipate the timing of rewards.

Anticipatory lick rates were slightly lower in the scanner (**Figures 2K**), but this was likely due to improvements in their learning of the task epochs, rather than excessive stress, as the lick rate to the outcome remained high (**Figures 2M**). Importantly, the difference between CS+ and CS- licking was still reliable (normalized lick rate: CS+: 2.4 ± 1.28, CS-: 0.43 ± 0.26, mean ± SD, t(4) = 3.23, p < 0.05, N = 5 mice, **Figure 2I**).

### Lick-related movements are coupled to signal artefacts that can be corrected

To measure brain-wide activity, we recorded changes in blood-oxygenation-level-dependent (BOLD) fMRI signals across the brain. Functional images (0.2 x 0.2 x 0.75 mm) were acquired at 1Hz with minimal geometrical distortions after optimising the setup, and the surgical and imaging procedures (**Figure 3A**). The head-fixation and training procedures were effective in minimising head-motion, as evidenced by the minimal head displacement observed across frames (mean framewise displacement across mice: 0.0081 ± 0.0012 mm, mean ± SD, N = 5 mice, note that these values are < 5% of the voxel size (0.2mm), **Figure S1**). However, previous fMRI studies in rats (Tabuchi et al., 2002), monkeys (Keliris et al., 2007) and humans (Van de Moortele et al., 2002) have shown that even with minimal changes in head position, detrimental image artefacts can be indirectly induced by movement of other body parts. Although previous fMRI studies in behaving mice did not investigate or report such effects (Han et al., 2019; Sakurai et al., 2020), we expected that similar artefacts would be also found in the mouse. Indeed, despite minimal brain movement, we found large amplitude changes in brain signals (**Figure 3D**) that were temporally coupled to lick events (**Figure 3B**), which require jaw and tongue movements. The near-global nature of this signal changes, indicated by the vertical lines in **Figure 3D**, is evidence that these changes are artefactual and not true fluctuations in brain activity. We exploited the fact that our field of view included large parts of muscle tissue (including the jaw and the tongue), and that our measurement is affected by geometrical changes in those voxels, to investigate whether indeed these artefacts could be arising from movements in those regions (e.g. muscle contractions and tongue movements). Indeed, video inspection revealed large geometrical changes in nearby muscle tissue, which when plotted over time (**Figure 3C**) revealed to be tightly coupled to the artefactual signal changes seen in the brain (**Figure 3D**). Note the consistency between the vertical lines in **Figures 3C** and **3D**, and the similarity between the global signal traces (averaged across voxels) in **Figure 3F**.

**Figure 3.**
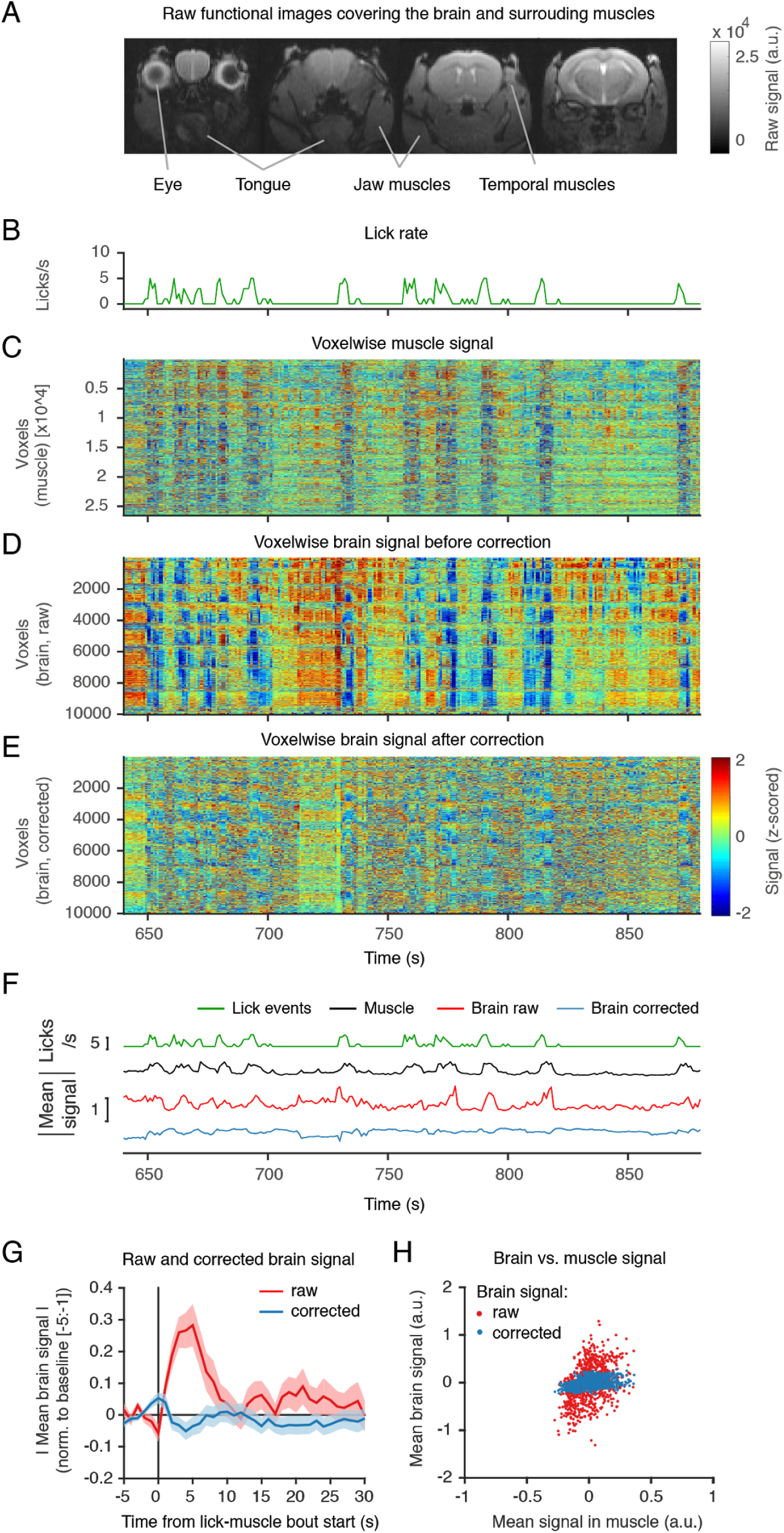
Lick-related muscle movements are coupled to signal artefacts that can be corrected using a muscle-drive regression approach. **(A)** Example raw functional images (0.2 x 0.2 x 0.75 mm at 1Hz) showing the brain and surrounding muscles. **(B)** Lick rate during imaging for a representative mouse. Shown is a zoom-in on a 250 s period. **(C)** Voxelwise time course for all “muscle” voxels (voxels outside the brain) for the same example period. Images were slice-timing and motion corrected, and the time course of each voxel detrended and z-scored. Note the vertical lines corresponding to geometrical changes (contractions) in the muscle tissue surrounding the skull. **(D)** Same as (C) but for voxels inside the brain. Large amplitude changes in the brain’s signal (vertical stripes) are temporally coupled to lick events (B) and muscle changes (C). **(E)** Same as (D) but after applying the artefact correction procedure. We used least absolute shrinkage and selection operator (LASSO) regression to predict the signal artefact in the brain from the information in the muscles voxels. After standard MRI preprocessing procedures, we created two binary masks of brain-only and non-brain voxels. We then computed the contribution of each voxel outside the brain at time i, to the intensity of each voxel in the brain at the same time i. To account for the fact that licks may fall in some but not all slices within the same frame, we treated slices independently. We then subtracted the artefact prediction from the brain signal to generate artefact corrected images (see Methods). **(F)** Time course comparison for lick events (licks/s), muscle signal (averaged over voxels), raw brain signal (averaged over voxels), and corrected brain signal (averaged over voxels). Shown are absolute values. Note the similarities between the upper 3 traces and the improvement after correction in the bottom trace. **(G)** Averaged time course of brain signal (raw or corrected) aligned on the start of a lick bout. The time courses (absolute values) were first averaged over voxels, bouts of consecutive licks were identified, time courses were aligned to the beginning of each bout, normalized (signal-baseline/baseline) to the mean signal before the bout started (-5:-1s from bout start) and finally averaged over bouts. Error bars represented SEM over all bouts in the session. **(H)** Scatter plots relating the brain and muscle mean signals for raw (red) and corrected (blue) brain signals. Each dot is a time point over the entire imaging session (890 frames).

It is unlikely that these artefacts are specific to our set-up as similar effects have been reported in monkeys (Keliris et al., 2007) and in the earlier study imaging rats while drinking water (Tabuchi et al., 2002). Behavioural strategies such as training animals not to lick during task-relevant periods are possible and have been used in the monkey literature (Keliris et al., 2007), however, they are significantly harder to achieve in rodents and highly restrict task designs. We thus aimed to tackle these artefacts not at the behavioural, but at the analysis level.

To do this, we exploited the rich information present in the muscle tissue to predict and remove the artefact in the brain. We estimated regression coefficients using least absolute shrinkage and selection operator (LASSO) (Tibshirani, 1996), as we had a large number of predictor voxels. LASSO is a regularisation method that, by pushing the coefficients of predictors (voxels in our case) that are least predictive to zero, effectively performs covariate selection, using only a subset of the regressors in the final model.

We preferred this approach over alternatives such as independent component analysis (ICA) as preliminary analyses revealed that there was not a clear separation of “lick” events (data not shown). This is likely because licks can manifest in many different ways (i.e. a first lick requires opening the mouth whereas a later lick in a bout occurs when the tongue is already fully extended) and are thus absorbed into several independent components. Because licks are highly correlated with task events (e.g. reward and odour), it is therefore hard to dissociate which components are related to which events.

Instead, by using the voxelwise signal over multiple muscles (i.e. modulated by muscle contraction), we gained access to the various possible configurations of the muscles and thus the various “types” of licking. This approach proved highly effective at minimizing these artefacts, evidenced by the: (1) reduced amplitude changes in voxelwise brain signal after correction (**Figure 3E**), (2) reduced global brain response upon lick-muscle movement (compare the red and blue trace in **Figures 3F** and **3G,** see also **Figure S2C** for cross-animal comparison), and **(3)** minimal linear correlation between muscle and global signal after correction (**Figure 3H,** see also **Figure S2D** for cross-animal comparison).

It is worth noting that changes in head muscle signal were also observed if mice were required to lever press in a separate task (**Figure S2B**). These changes were also coupled to brain signal artefacts that could be corrected using our muscle-driven approach (**Figure S2C** and **S2E**). This demonstrates one additional advantage of using the muscle tissue directly as the driver of artefact corrections: artefacts arising from any movements that cause contractions of these muscles will be corrected for, even if these movements did not lead to event detection (i.e. lick spout vibration or lever deflection).

Given the correlation between the artefacts and task-related movements, effective artefact identification and correction methods are crucial steps to achieve valid and interpretable results. Our results show that a muscle-based preprocessing strategy is effective at correcting these artefacts, without the need for additional behavioural training (Keliris et al., 2007) or the loss of large fractions of critical data by discarding affected frames. We next turned to the resulting brain activity for further validation of its effectiveness.

### Distinct spatial patterns for different odours in the olfactory bulb

We explored the rich activity patterns recorded in the olfactory classical conditioning task **(Figure 2B)**, beginning by looking for previously-described responses to validate our method. The main olfactory bulb (OB) receives input from the olfactory epithelium via olfactory receptor neurons (ORN). Each ORN expresses a single odourant receptor gene and ORNs of the same type converge to the same glomeruli in the OB. This means that in the OB, odourant receptors are arrayed into a spatial map of glomeruli, such that each odour activates a unique spatial pattern of activity (Korsching, 2001; Leon & Johnson, 2003; Mori, 2003; Rubin & Katz, 1999; Vassar et al., 1994). We thus first focused on the OB (**Figures 4A** and **4B**), where we expected to see distinct spatial patterns for different odours.

**Figure 4.**
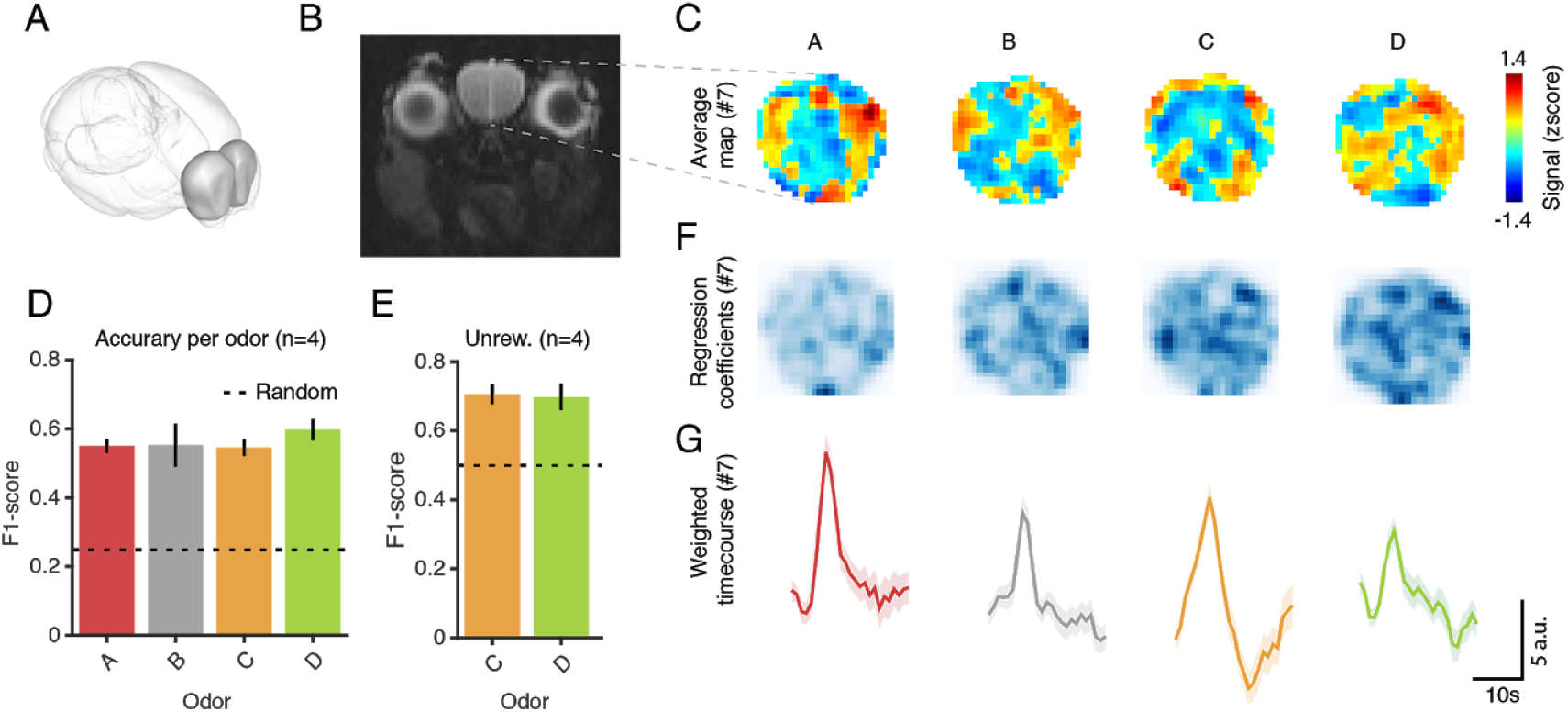
Odor identification in a classical conditioning task. **(A)** Allen brain atlas showing the olfactory bulb (OB). **(B)** fMRI coronal slice showing the OB (surrounded by the eyes, the tongue, and other head muscles). **(C)** Averaged maps of BOLD-fMRI signal for each odours for an individual mouse (#7). Voxelwise time courses were aligned on odour onset, pooled over trials of the same type, and averaged over a 10s window to account for the slow dynamics of BOLD. Colors indicate z-scores. **(D)** Decoding accuracy for odour identity considering the 4 odours. F1-score is a measure of accuracy, defined as: 2 * (precision * recall)/(precision + recall). Decoding was done for individual animals using multinomial logistic regression (a generalised linear classifier) in the same time window as (C), cross validating with stratified 5-fold. Unsmoothed images were used. Random accuracy is 0.25. **(E)** Decoding for unrewarded odours only, using the same approach. Random accuracy is 0.5 in this case. **(F)** Maps of regression coefficient for the same animal shown in (C), darker colours indicate voxels that contributed most to the decoding. **(G)** Time course of olfactory bulb activation weighted by the regression coefficients to reflect voxels most relevant for each odour’s discrimination. One animal was excluded from this and any further analysis as he showed poor task performance during testing.

We restricted our analysis to within-subject comparisons for two methodological reasons. First, given the small size of glomeruli (∼0.08 mm) (Royet et al., 1988), it was difficult to ensure that the same arrays of glomeruli, sampled in only 1-2 coronal slices of our whole-brain dataset, were imaged across mice. Second, given that we counterbalanced odour-reward contingencies across animals and that reward associations are known to also modulate the OB (Doucette et al., 2011; Kay & Laurent, 1999) we introduced cross-animal variability not related to the sensory representation. We thus looked for reproducible odour maps across trials for each subject while allowing for cross-animal variability.

Indeed, averaging maps of the OB over trials of the same odour revealed distinct spatial patterns for different odours (**Figure 4C,** see also **Figure S3**). To confirm that these patterns were reliable across trials for all subjects, we asked if a linear classifier could correctly identify the odour presented based on the trial-by-trial activity in the OB. We computed decoding accuracy per odour for each mouse before averaging across animals. The classifier performed above chance for all four odours (**Figure 4D)**. To ensure that this was not due to asymmetries in licking behaviour (in the two rewarded versus two neutral odours), we then restricted the decoding to unrewarded trials only. The performance was still above chance (**Figure 4E**), confirming the discriminability of different odours. **Figure 4F** shows the contribution of each voxel to the decoding and the corresponding averaged time courses over voxels weighted by these contribution coefficients (**Figure 4G**). As expected for the slow dynamics of BOLD-fMRI signals, responses evolved over several seconds upon odour delivery. These results are additional evidence for the effectiveness of our correction strategies.

### Motor and reward correlates in the odour-guided classical conditioning task

To take full advantage of this classical conditioning task, we next explored how brain-wide odour responses depended on their reward contingencies. Based on the literature we expected the olfactory bulb and the olfactory (piriform) cortex to show reward related signals (Calu et al., 2007; Doucette et al., 2011; Kay & Laurent, 1999; Roesch et al., 2006). Beyond the olfactory system, we expected that reward and neutral odours would be followed by differential activity in both reward-related areas, associated with the different reward contingencies, and motor-related regions involved in licking behaviour.

To do this, we ran a voxel-wise general linear model (GLM) with CS+, CS-, US+, US- as covariates (Friston et al., 2007). We then contrasted rewarded and unrewarded conditions. The resulting statistical maps are shown in **Figures 5A** and **5B**. In the odour (CS) period, we found voxels within the olfactory bulb and the primary (piriform) cortex (**Figure 5A**). In the outcome (US) period, we found multiple cortical and subcortical areas (**Figure 5B**). As expected, we found engagement of multiple reward-related areas such as the amygdala (AMG) and the anterior cingulate area (ACA), known to encode value (Cardinal et al., 2002; Janak & Tye, 2015; Kolling et al., 2016), as well as multiple hypothalamic areas (lateral hypothalamic area, LHA; medial/median preoptic area, MPO, and medial preoptic nucleus, MPN; paraventricular hypothalamic nucleus, PVH). In particular, the medial/median preoptic area has been implicated in thirst regulation and drinking behaviour (Abbott et al., 2016; Ji et al., 2005; McKinley et al., 1994).

**Figure 5.**
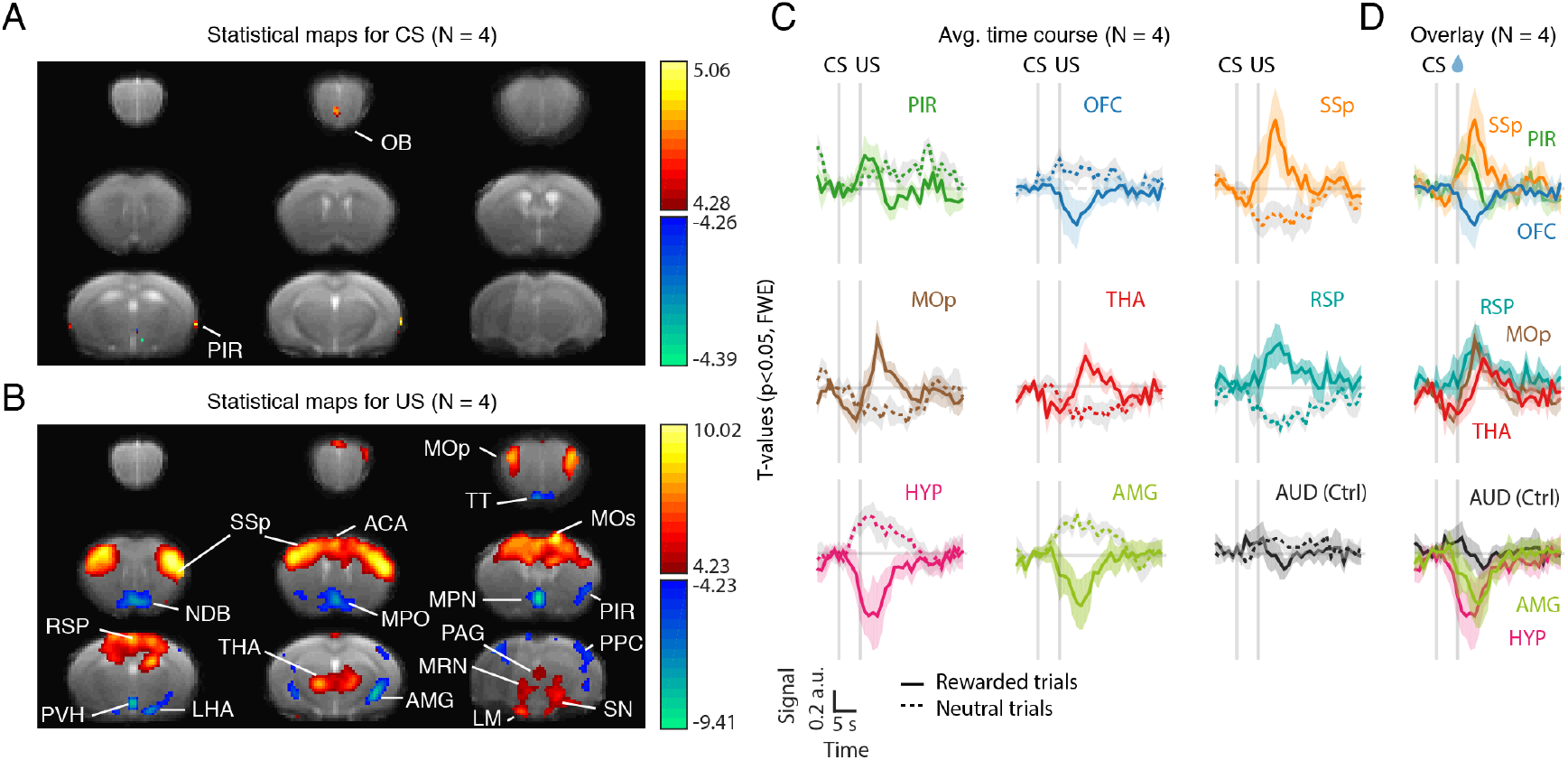
Motor and reward correlates in the classical conditioning task. **(A)** T-statistical maps (N = 4 mice) for the contrast between the CS+ and CS- regressors. Thresholded at p<0.05 and corrected for multiple comparisons with Family-Wise Error (FWE) correction. Hot colors correspond to areas more active in the CS+ condition than in the CS-. Cold colors correspond to areas less active in the CS+ condition than in the CS-. Data pooling 4 mice. One animal was excluded on the basis of poor task performance (little licking behaviour) during testing. **(B)** Same as (A) but for the US+ and US- regressors. **(B)** Averaged time-course in regions-of-interest (ROIs) identified in the GLM analysis, aligned on trial start (odour onset). After standard preprocessing (slice timing, motion correction, detrending) and artefact correction, images were z-scored voxel-by-voxel. For each ROI, z-scored time courses were averaged over voxels, aligned on odour onset, baseline subtracted (average of -2:-1 s prior to odour onset), averaged over trials for either rewarded trials (solid line) or unrewarded trials (dotted line) and, finally, averaged across mice. Error bars represent SEM across mice. **(C)** Overlay of the time courses shown in (B) for rewarded trials only. Note the difference in onset and peak response across areas. CS: Conditioned stimulus. US: Unconditioned stimulus. Ctrl: control region. For abbreviations of brain regions see Table S1.

Consistent with our predictions, we also found multiple motor-related regions (**Figure 5B**) including the midbrain reticular nucleus (MRN), primary and secondary motor cortices (MOp, MOs), thalamus (THA), and somatosensory cortex (SSp). Given our spatial resolution and the spatial blurring inherent to haemodynamic measures, we cannot distinguish substantia nigra reticulata (motor-related) from its reward-related neighbours (ventral tegmental areas and substantia nigra pars compacta). We thus termed them conservatively as SN. Additional areas identified in our analysis included the taenia tecta (TT), which receives sensory input from the olfactory bulb and top-down input from prefrontal cortex (Hoover & Vertes, 2011; Igarashi et al., 2012), the retrosplenial cortex (RSP), implicated in learning and memory (Vann et al., 2009), the diagonal band nucleus (NDB), part of the basal forebrain and source of cholinergic projections (Mesulam et al., 1983), mammillary nucleus (LM), implicated in memory (Vann & Aggleton, 2004), periaqueductal gray (PAG), implicated in autonomic regulation and the expression of both aversive and appetitive responses (Motta et al., 2017; Tryon & Mizumori, 2018), posterior parietal cortex (PPC), implicated, among other functions, in the planning and control of movement (Andersen & Cui, 2009; Y. E. Cohen & Andersen, 2002).

To further investigate the activity of these areas we plotted and compared their time courses aligned on odour onset (**Figure 5C** and **5D**). As expected, we saw activation of olfactory areas such as piriform rising before outcome delivery and, preceding activation of motor lick-related areas such as primary somatosensory cortex (**Figures 5D**, top panel). Most areas peaked after reward delivery (**Figure 5D**). However, given the delayed nature of the BOLD response in relation to its underlying neural activity, this is expected even in the case of areas that were activated prior to the outcome. Nonetheless, in some cases, including MOp and THA, changes appeared even before the US presentation (**Figure 5D**), consistent with the expression of anticipatory licking. Interestingly, responses in the HYP also started before reward delivery and prior to other areas such as the AMG (**Figure 5D**, bottom), suggesting that it may be involved in the anticipation, in addition to the consumption of water reward. Consistent with this hypothesis, a recent study found that optogenetic activation of neurons in the medial, also known as median, preoptic nucleus of the hypothalamus promotes operant lever pressing for water rewards at rates that scale with stimulation frequency (Allen et al., 2017). As a control region, we looked at the auditory cortex (AUD). By contrast to the other areas, AUD showed little modulation (**Figure 5C**, last panel).

Taken together, these results offer strong validation for our method, highlighting a number of areas that have been traditionally studied in isolation and individually implicated in the sensory, reward and motor processes required for this task.

### A self-paced operant conditioning task

We next aimed to increase complexity and degrees of freedom to the behavioural task by adding a preparatory action (lever pressing) that was distinguishable from the action needed for reward consumption (licking). Contrary to licking, which is reflexive, lever is learnt and voluntary, and thus offers a separate and more controlled readout for the study of cognitive phenomena such as decision-making. Our motivation was therefore: first, to make our approach compatible with a broad range of decision-making tasks that can separate, at the behavioural and neural level, preparatory behaviour or choice from reward consumption. Second, by having multiple behavioural outputs, including a right and left lever pressing, we aimed to make our approach compatible with the use of more complex, quantitative paradigms such as 2-alternative forced-choice tasks, where behaviour can be carefully modeled. Finally, we aimed to explore the neural correlates of learned, self-paced preparatory actions. To do this, we chose a self-paced task and imaged over a learning period so that the variability in the behaviour, both between individual presses and between presses and licking, was maximised and thus helped parse their neural correlates.

Using the same setup and training procedure described in **Figures 1** and **2**, we trained mice to press one of two levers in the scanner to obtain delayed rewards. In this task, pressing a lever (either left or right) with the forepaw, in the appropriate period, resulted in the delivery of a water reward available upon licking after a delay (**Figure 6A**). During the early stages of training, every single press (left or right) was reinforced with a water reward. This produces reliable pressing but meant that mice would become satiated quickly and that licks were tightly coupled with presses. To slow down the behaviour and start decoupling presses from rewards, and thus licking, we inserted a cued waiting period (2-4s, increased over sessions) during which presses were not rewarded. The first press after the waiting period elapsed was termed *valid press* and was rewarded after a delay (0.5 - 2s, increased over sessions). A blue light provided immediate positive feedback after a valid press and stayed on until reward delivery, serving also as a bridging cue. Reward delivery was followed by a short inter-trial interval (ITI, 0.5s) (**Figure 6A**).

**Figure 6.**
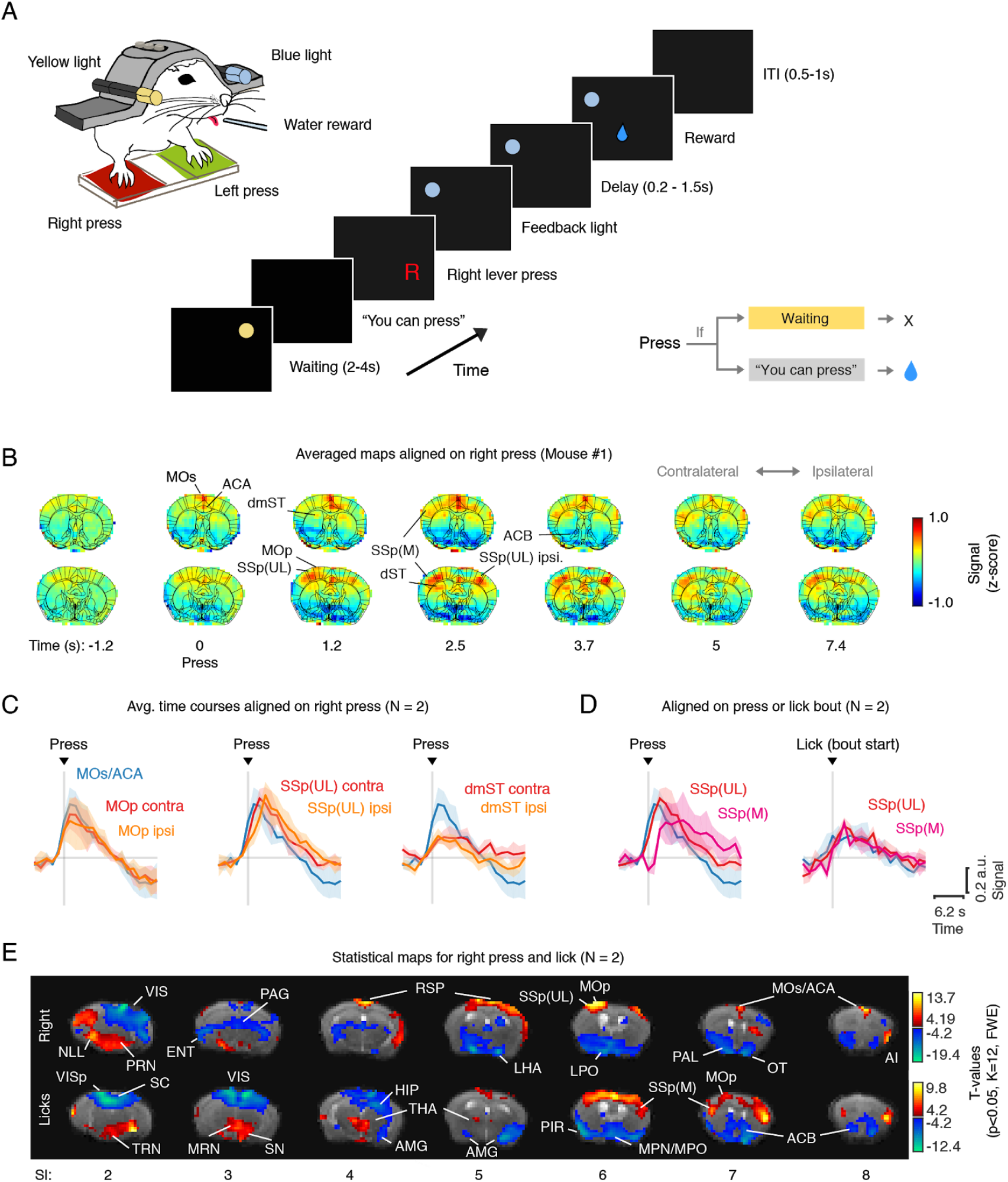
Brain-wide task correlates in a self-paced operant conditioning task. **(A)** Schematic of the lever press setup (left) and operant conditioning task structure (centre). A trial started with a yellow cue in the right visual field signaling the waiting period (2-4s, increased over sessions). Presses during this period were neither rewarded nor punished. The first press (either right or left) after the waiting period had elapsed, was defined as a valid press and was rewarded after a delay (0.5-1.5s, increased across sessions). A blue light to the left of the animal provided immediate positive feedback after a valid press and stayed on until reward delivery. Reward delivery was followed by a short inter-trial interval (0.5s). Differences in outcome between the two press types (valid and invalid) (right). **(B)** Spatiotemporal maps over two coronal slices aligned on right lever pressing for a single animal (mouse #1). Event selection was further constrained to exclude right presses preceded or superseded by left presses in a window of [-10:10]s. After standard preprocessing (slice timing, motion correction, detrending) and artefact correction, images were z-scored voxel-by-voxel, aligned on right press events, baseline subtracted (average map in -2:-1s from press), and averaged over events. Colors indicate normalized z-score signal. **(C)** ROI time courses aligned on right lever presses, averaged over mice (N = 2). Error bars represent SEM across animals. **(D)** Comparison of primary somatosensory time courses in the upper-limb and mouth regions, aligned on either right lever press (left panel) or lick bout start (right panel). **(E)** T-statistical maps for the right and lick regressors pooling both animals. Thresholded at p<0.05, minimum cluster size (k) = 12, and corrected for multiple comparisons (FWE). For abbreviations, see Table S1.

The two mice that we trained and tested pressed several times over a session (121 ± 58 presses, 40 ±16 rewarded, mean ± SEM, from a total of N = 14 sessions from 2 mice), with a preference for right lever press (77 ± 14% across sessions). Consistent with our aim of maximising the variability in the behaviour, the interval between subsequent presses was highly variable, as indexed by the difference between the 90th and 10th percentiles of the inter-press intervals (IPI) (Delta IPI _[0.1 - 0.9]_ median = 60s, range = 17 - 196s).

### Motor and somatosensory correlates of right lever presses

We began by looking for press-related brain activity, which we expected to engage striatal, motor, and somatosensory regions, areas implicated in the planning, execution, and sensory feedback of movements (Kawai et al., 2015; Krakauer et al., 2019; Li et al., 2015; Luft & Buitrago, 2005). We focused on right presses as they accounted for the majority of lever presses. **Figure 6B** shows the spatiotemporal maps of fMRI-BOLD signals averaged across right presses and aligned on press detection for an individual animal. As predicted, we detected activation in MOp, MOs, dorsal striatum (dST) and the upper limb region of the somatosensory cortex (SSp(UL)). The activity started in the ACA and MOs, followed by activation of dST, MOp and the SSp(UL), contralateral to the press. Activity in the ipsilateral SSp(UL) appeared only later. These results were consistent across the two mice tested, as shown in the averaged time courses over these areas (**Figure 6C**). The asymmetry between hemispheres will be further explored in a later multivariate analysis where the contribution of left and licks can be better accounted for.

### Separating preparatory and consummatory activity

Next, we aimed to parse the neural correlates of preparatory (lever pressing) from consummatory (licking) behaviour. We started by focusing on the somatosensory cortex where we know that distinct body parts are represented in spatially distinct areas (Coq & Xerri, 1998). Licking behaviour should engage preferentially the mouth area, whereas lever pressing should engage the upper limb area. Indeed, aligning activity of these regions on lever press reveals BOLD-fMRI signal increases in the upper limb region at the time of press, followed by activity in the mouth region (SSp(M)), consistent with the subsequent consumption of the water reward (**Figure 6D**, left). Accordingly, aligning activity in the same areas to the start of a lick bout shows mouth area activity coupled to licking (**Figure 6D**, right).

To explore lever-press and lick-related activity across the brain, we ran a voxelwise GLM analysis. Using GLMs allowed us to exploit trial-by-trial variability in the behaviour, by including right lever presses, left lever presses and licks as covariates. Given the low number of left presses and the fact that mice often performed them simultaneously with right ones, we treated the left as a nuisance regressor. The resulting statistical maps are shown in **Figure 6E**. Consistent with our previous analysis, we found that right lever presses preferentially engaged the upper region of the somatosensory cortex, while licking correlates best with the mouth region. Furthermore, we found that although both lever press and licking engage motor and somatosensory area, lever press engages preferentially the contralateral hemisphere, whereas licking is more bilateral, as also seen in the classical conditioning task. The contralateral bias for right press is clearer in this analysis relative to the averages (**Figure 6B**) as the contributions of brain activity related to licking and left-presses can be better parsed from activity related to right-lever pressing.

In addition to these areas, we found task-related activity in several additional cortical and subcortical regions (**Figure 6E**). Among the areas best correlated with right lever press were retrosplenial cortex (RSP), insular cortex (AI), several midbrain motor nuclei (NLL, PRN), pallidum (PAL), olfactory tubercle (OT) and lateral hypothalamus (LHA). Licking engaged preferentially areas such as midbrain reticular nucleus (MRN), substantia nigra (SN), hippocampus (HIP), amygdala (AMG), piriform cortex (PIR), nucleus accumbens (ACB), parts of the thalamus (THA) and more medial aspects of the hypothalamus (MPN and MPO). Some of these areas, such as motor thalamus and hypothalamic areas overlap with the areas identified in the classical conditioning task, consistent with the use of comparable water reinforcement and the expression of licking. The AMG was also detected in both tasks. The basolateral AMG, in addition to playing a role in Pavlovian stimulus-outcome associations, has also been shown to use this information to modulate instrumental actions (Cardinal et al., 2002).

Finally, given the use of light cues in the task, we also expected to see the engagement of visual areas such as visual cortex (VIS) and superior colliculus (SC). Valid presses are immediately followed by a blue light (feedback signal) that is positioned on the left side of the visual field (**Figure 6A**). We thus expected them to correlate with contralateral (right) visual cortex activity. Licking, by contrast, occurred both during the reward period (cued by the left blue light) and during the initial part of the subsequent waiting period (cued by the right yellow light). We thus expected licking to engage visual areas bilaterally. Indeed, **Figure 6E** confirms these predictions.

### Using decoding approaches to further investigate task-related activity

To further investigate what information may be available in these areas, we took advantage of decoding strategies. We exploited the fact that we had a large dataset per mouse in this task, to use support-vector machine algorithms (Suykens & Vandewalle, 1999). We started by comparing valid (“correct” press, first press after the waiting period had elapsed) and invalid presses (presses during the waiting period) (**Figure 7A**). They should be comparable in terms of kinematics for pressing. Importantly, they differ in their reward contingencies: whereas a valid press is always followed by a delayed reward, invalid presses are never rewarded. In addition, they will also differ in visual stimuli: whereas invalid presses are correlated with the constant yellow light of the waiting period, presented in the right visual field, valid presses are immediately followed by the transient blue light, presented to the left of the visual field.

**Figure 7.**
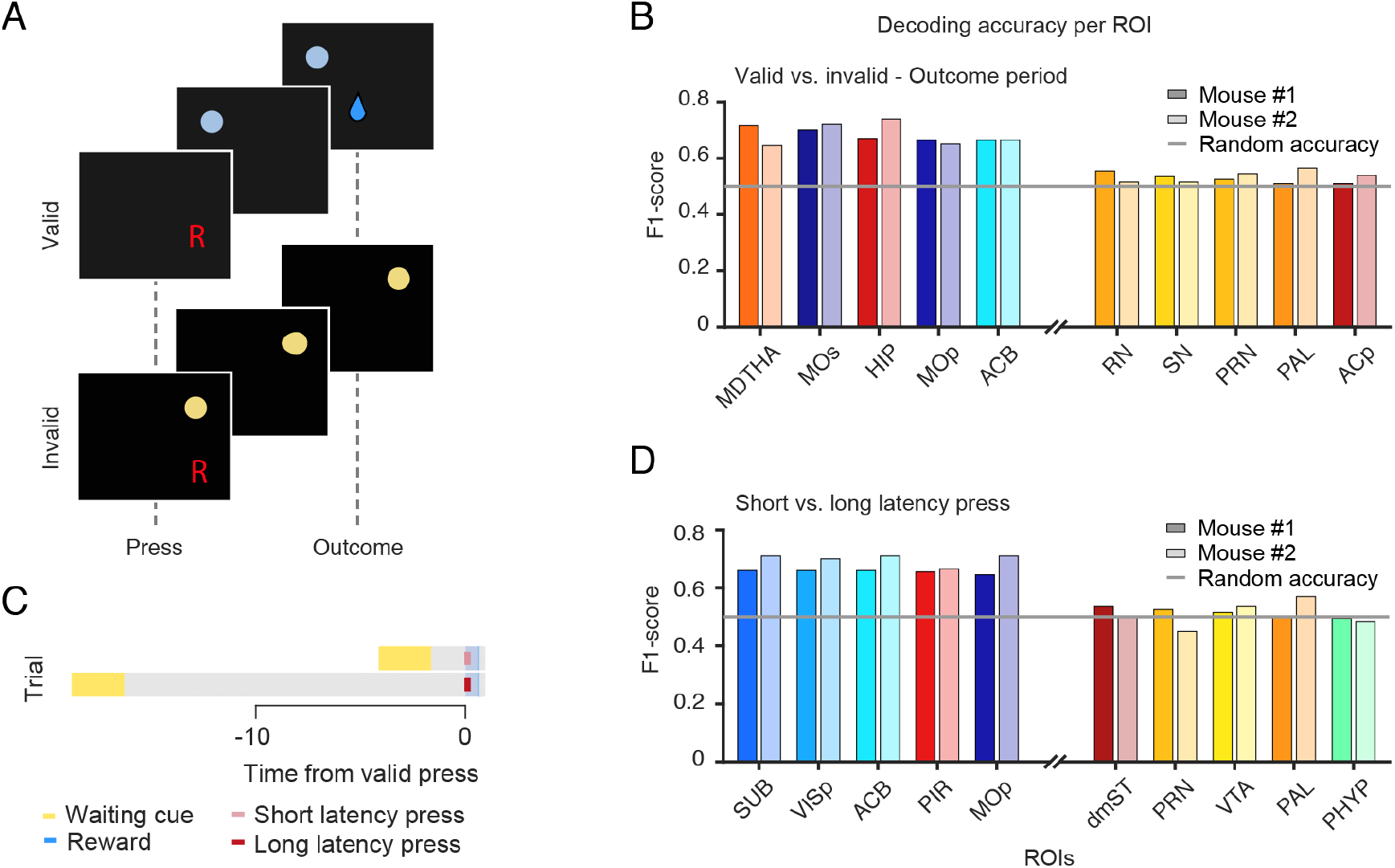
Decoding task-relevant activity from multiple areas. **(A)** Schematic of valid and invalid presses. Whereas valid presses are always followed by a delayed reward. Invalid presses are never rewarded. **(B)** Region-by-region accuracy in the decoding of valid versus invalid press during the outcome period (1:5s after press). F1-score is a measure of accuracy, defined as: 2 * (precision * recall)/(precision + recall). Decoding was done for individual mice pooling data from all sessions (N = 7) using Support Vector Machine algorithms, cross-validating with stratified 5-fold. Unsmoothed images were used. Areas were first ranked according to their accuracy. Among the 15 most and least predictive, we asked which are common across mice. Shown are the 5 most (least) predictive areas that were present in the top (bottom) 15 for both mice (see Methods for more details). Mouse #1 is in dark shades, Mouse #2 in light shades. Random accuracy is 0.5. **(C)** Schematic of short-latency and long-latency presses. The self-paced nature of the task meant that valid presses occurred after variable delays. We split latencies into a lower and upper percentile, defining trials with latencies below the 50th percentile as short and those above as long. **(D)** Region-by-region accuracy in the decoding of short vs. long latency presses in a period around the time of press (-3:3s relative to press).

Multivoxel decoding analysis during the outcome period (1:5s after press) revealed that among the top areas to discriminate the two conditions were mediodorsal thalamus (MDTHA), secondary motor cortex (MOs), hippocampus (HIP), primary motor cortex (MOp) and nucleus accumbens (ACB) (**Figure 7B**). The MDTHA is a higher-order thalamic nucleus that receives inputs from medial prefrontal cortex, limbic structures and basal ganglia (Marton et al., 2018). Its function is not well understood but mounting evidence suggests a role for MDTHA in associative learning (Bradfield et al., 2013; Chakraborty et al., 2016; Izquierdo & Murray, 2010; Mair et al., 2015; Mitchell, 2015; Mitchell & Chakraborty, 2013; Ostlund & Balleine, 2008).

MOs is involved in action planning and initiation (Erlich et al., 2011; Guo, Li, et al., 2014). It has also been shown to convey value signals before and after a choice has been made (Sul et al., 2011). Its involvement in our task is likely to reflect both the planning/initiation of consummatory actions (i.e. licking to collect reward) and the outcome difference between valid and invalid presses. Consistent with a role in action planning and initiation, primary motor cortex (MOp) is also among the most predictive areas. In line with a role in value coding, the ACB, known to be involved in reward processing (Cardinal et al., 2002), was also among the most predictive areas.

The HIP has been implicated in learning and memory (Eichenbaum et al., 1999; Good, 2002; McNaughton & Morris, 1987; Morris & Frey, 1997). In the context of reward processing, it has been shown to receive input from reward-related areas to enhance encoding of rewarding events (Adcock et al., 2006; Lisman & Grace, 2005; Murty & Adcock, 2014; Shohamy & Adcock, 2010; Wittmann et al., 2005). Reward-related differences in memory encoding could underlie the hippocampal contribution to discriminating the reward and no-reward conditions in our task.

By varying in outcome, valid and invalid presses will also vary in the extent to which they engage licking. We next compared conditions that were matched not only for press output but also for licking: short- and long-latency valid presses (**Figure 7C**). The self-paced nature of our task meant that mice could press at any point after the waiting period had elapsed. We defined this interval (time from the end of the waiting period to the first press) as the “latency to press”. We then split presses into short-latency (<50th percentile of all latencies) and long-latency (>50th percentile). Because the first press after the waiting period (whether short or long-latency) was always rewarded, these conditions are comparable in both press and lick-related movements, but they should differ in terms of the decision, motivation and preparation involved in making an early, as opposed to a delayed, action. Again, since short-latency presses happen temporally closer to the end of the waiting period, these conditions also differ in visual stimuli.

Among the areas best at decoding short- from long-latency presses were subiculum (SUB), primary visual cortex (VISp), nucleus accumbens (ACB), piriform cortex (PIR), and primary motor cortex (MOp) (**Figure 7D**).

In our task, short- and long-latency presses may imply different motivational states that modulate the latency to act. Consistent with this hypothesis, both the SUB and ACB have been implicated in motivated reward-seeking behaviour (Cardinal et al., 2002; Vorel, 2001). In particular, stimulation of SUB has been shown to be effective in eliciting reinstatement of cocaine-reward seeking when delivered before lever pressing, suggesting that it has predictive or incentive properties that facilitate action initiation (Vorel, 2001). The ACB works at the interface between reward and motor systems by allowing motivational stimuli to enhance ongoing instrumental responding (Pavlovian-instrumental transfer, PIT) (Cardinal et al., 2002). Although PIT is commonly measured in terms of response vigor, it seems possible that it would also be expressed in terms of the latency to press. Interestingly, stimulation of SUB induces long-lasting dopamine (DA) release in ACB (Blaha et al., 1997; Brudzynski & Gibson, 1997; Legault et al., 2000; Legault & Wise, 1999), and locomotion towards a goal alters synchronous firing of neurons recorded simultaneously in SUB and ACB (Martin, 2001).

Note that given that short-latency presses are temporally closer to the reward in the preceding trial compared with long-latency presses, we cannot rule out that some of the activity of the previous reward contributes to this decoding. This could underlie at least partly the ACB and MOp also identified in the decoding of the outcome. However, given the differential involvement of other areas, it is also possible that they play both, potentially related, roles.

PIR is the largest of the olfactory cortical areas and has been shown to be involved in processing olfactory information (Haberly, 2001), both in terms of its sensory and associative features (Calu et al., 2007; Roesch et al., 2006). No study to our knowledge has investigated piriform in the context of non-olfactory tasks. Our findings raise the interesting possibility that it may also play a role in non-olfactory associative processes. Further studies are however needed to verify this preliminary conclusion.

Together these findings illustrate how whole-brain fMRI data, combined with advanced analysis techniques, can be used to aid unbiased investigation of the brain-wide circuits mediating behaviour. By providing simultaneous information across multiple brain regions active during behaviour, this information can be used to form hypotheses that can then be used using complementary invasive circuit techniques.

### Using brain-wide correlational analysis to investigate relationship between areas

Behaviour depends not only on intra-area local processing but critically on the interaction and coordination of information processing across multiple areas. We exploited the ability of fMRI to provide information about multiple areas simultaneously recording across the whole-brain to explore functional coupling between them. We define functional coupling as the overall correlation between voxel or ROI time series. This will include, besides functional connectivity, any coherence in task-driven responses. We first focused on the ACB, a well-studied subcortical region, known to work at the interface between reward and motor systems by allowing motivational stimuli to shape instrumental responding (Cardinal et al., 2002). We explored functional coupling by computing the correlation between the mean time course of the selected (seed) area (ACB) with every other voxel in the brain. **Figure 8A** shows the resulting maps for the two mice tested. We found functional coupling with regions including the amygdala, midbrain structures, motor and reward related structures (MRN, SN/VTA), hypothalamus, as well as motor and somatosensory regions, a pattern broadly consistent with the known function of ACB. Interestingly, we also identified the hippocampus, which we previously identified as predictive of the outcome, along with ACB.

**Figure 8.**
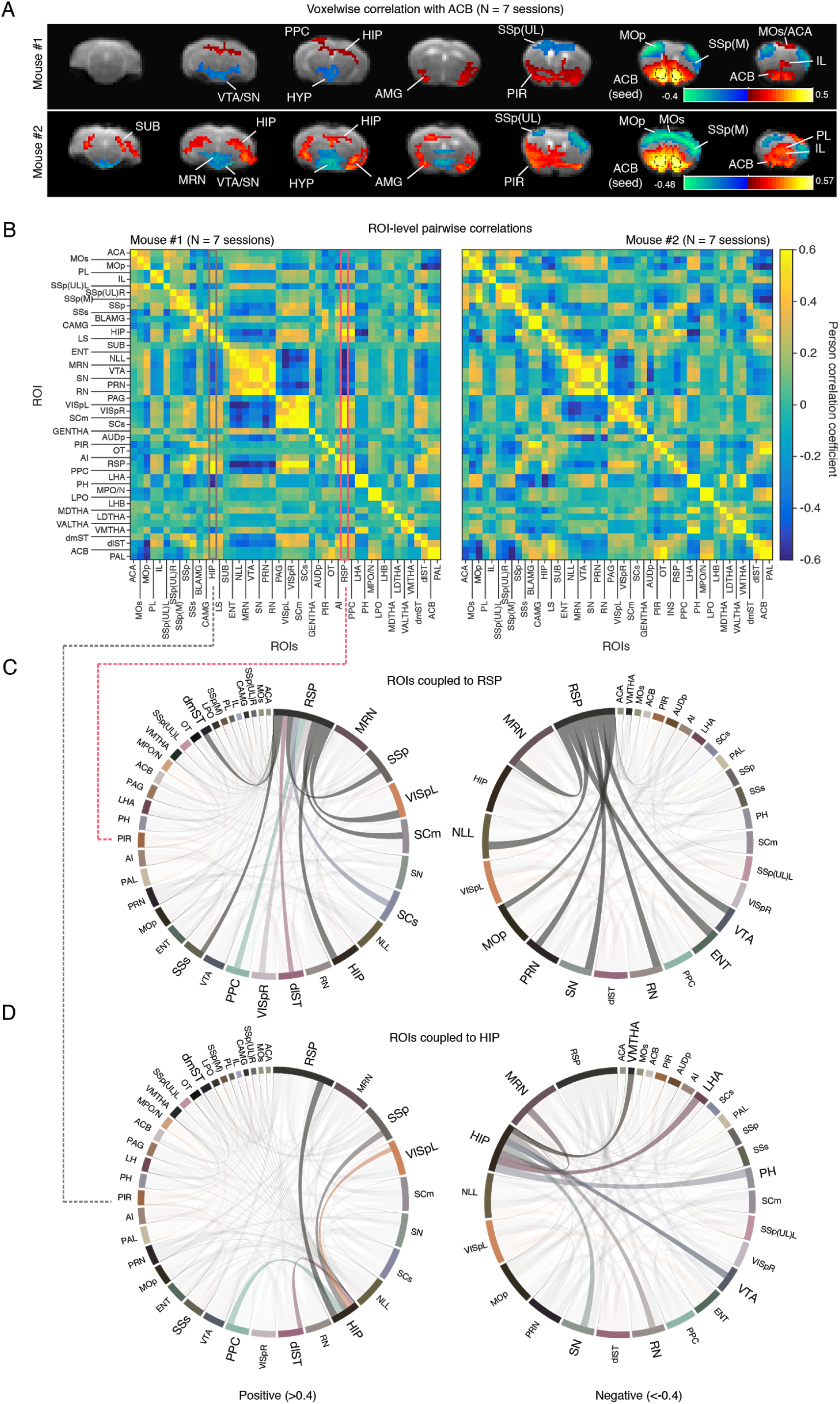
Functional coupling between multiple brain areas. **(A)** Seed analysis. Voxelwise correlation with seed region (nucleus accumbens, ACB). Columns are different coronal slices from posterior (left) to anterior (right). Rows are different mice. Colours are positive and negative Pearson correlation coefficients, thresholded at +- 0.2. **(B)** Pairwise Pearson correlation coefficients between various regions-of-interest (ROIs). Coefficients were calculated per session and averaged across sessions (N = 7). **(C)** Chord diagram showing correlations with retrosplenial cortex (RSP) thresholded at 0.4. The thickness of the line indicates strength of correlation. Positive correlations of the left. Negative correlations on the right. **(D)** Chord diagram showing correlations with hippocampus (HIP) thresholded at 0.4. Positive correlations of the left. Negative correlations on the right. The interactive version of these chord diagrams is available at https://madalena_fonseca.gitlab.io/coupling_visualisation/.

We next aimed to explore the functional coupling of some of the areas identified in previous analyses. We thus quantified the pairwise correlation among all voxels (**Figure S4**) or across ROIs (**Figure 8B**). To better visualise this information, we plotted it as an interactive chord diagram where the size of the connections between areas reflects the strength of their correlation. We first looked at the retrosplenial cortex (RSP) (**Figure 8C**), which we identified in the GLM analysis, most strongly correlated with the right press predictor, suggesting it may be more involved in the appetitive behaviour rather than reward consumption. RSP is at the intersection of areas that encode visual information, motor feedback, higher-order decision-making, and the hippocampal formation (Kononenko & Witter, 2012; Miyashita & Rockland, 2007; Sugar et al., 2011; van Groen & Wyss, 1992). Previous studies have implicated it in spatial navigation guided by visual or self-motion cues (Elduayen & Save, 2014), as well as learning and memory (Vann et al., 2009). Interestingly, within the context of its role in spatial navigation, it has been suggested that RSP may specifically contribute to cognitive processing by integrating visual, and potentially motivational, cues with information generated by self-motion (Cooper & Mizumori, 2001). This raises the interesting possibility that although not in the context of space, RSP may be involved in our operant task in using visual cues, and potentially their reward association, to guide, and learn from, instrumental actions.

Consistent with its relationship to the visual and hippocampal systems, among the higher correlations are visual cortex (VISp), hippocampus (HIP), and entorhinal cortex (ENT). Additional areas include motor cortex (MOp), and both dorsomedial (dmST) and dorsolateral striatum (dlST), consistent with its involvement at the time of pressing. Perhaps surprisingly, supplemental somatosensory cortex (SSs) was also among its higher correlations.

The role of SSs in behaviour is poorly understood but there is some evidence for its involvement in processing higher-order features of somatosensory stimuli including attention (Burton, 1999; Chapman & Meftah, 2005; Fujiwara et al., 2002; Meftah et al., 2002; Mima et al., 1998), learning (Debowska et al., 2011; Manzoni et al., 1979; Quallo et al., 2009), and sensorimotor integration (Huttunen et al., 1996). No studies to our knowledge have looked at SSs in the context of an operant task. Consistent with a role in attention and learning, SSs could be involved in our task in learning from tactile sensory and reward feedback to guide appropriate forelimb lever press movements.

To verify the robustness of our functional coupling analysis to spatial smearing of the BOLD signal, we compared the functional coupling of RSP to that of an adjacent area, the hippocampus (HIP) (**Figure 8D**), which we identified in the decoding analysis as discriminating reward outcome. Given their different potential involvements in the task, we expected to see differences in functional coupling. As expected from being correlated with each other, HIP shared correlation with RSP. These included VISp, posterior parietal cortex (PPC), dlST, and some midbrain motor and reward-related nuclei (RN, SN/VTA, MRN). However, while RSP showed a high correlation with MOp, SSs, and ENT, HIP showed a high correlation with primary, rather than supplemental, somatosensory cortex (SSp), lateral and posterior hypothalamic areas (LHA, PH), and ventromedial thalamus (VMTHA), consistent with the finding from the decoding analysis discriminating behaviour at the time of outcome delivery.

By using brain-wide correlation analysis, we exploited functional coupling between regions to further investigate the role that different regions may play in our task. We compared retrosplenial cortex and hippocampus, two areas that we had previously identified. Although in our case, the use of two different behavioural actions (lever and press) already allowed us at the GLM level to somewhat narrow their contributions, functional coupling provides a way to access this information potentially independent of behavioural labels or expression. For instance, one can imagine performing this analysis comparing behaviourally relevant conditions (e.g. before and after learning, or during different delay periods) that may not lead to differences in expressed behaviour but may reflect covert differences in cognitive processes. We explored one variant of this analysis, but there are several possible extensions, including how functional coupling changes over time (e.g. over learning) and how it relates to functional connectivity (e.g. by removing task-related activation), potentially providing a new layer of information.

## DISCUSSION

Access to a brainwide map of simultaneously recorded brain activity during mouse behaviour would inform system-levels theories of how dynamic distributed processing relates to behaviour and offer an unbiased way to select brain regions to be studied at the local level. We present a number of hardware, procedural, and analysis advances to enable the use of mouse behaving fMRI to tackle a broad range of behavioural neuroscience questions. First, we make MRI compatible behavioural setups compatible with the use of multiple sensory stimuli and behavioural readout measures, key to dissociate sensory, cognitive, and motor processing. Second, we identify and correct important signal artefacts from task-related body movements not addressed by previous studies. Third, we validate our approach in a 4-odour guided classical conditioning task by showing engagement of a number of regions previously implicated in classical conditioning as well as others that have been less studied. Finally, we extend our approach to a more general, operant task, with increased motor complexity and design flexibility. We further illustrate how a combination of analysis tools can be exploited to identify and investigate task-related activity across multiple cortical and subcortical areas and to explore brain-wide inter-region functional coupling.

The study of neural activity patterns during complex behaviour in rodents is essential to understand the mechanisms underlying animal behaviour and cognition. However, exactly to what extent findings in rodents relate to those in humans is far from clear. This comparison has been hampered by the fact that rodent and human studies use typically different techniques, measure different types of signal, and study the brain at drastically different spatial and time scales. Behaving rodent fMRI offers the possibility of measuring the same signals in comparable behavioural tasks. This data can then be used not only to find similarities but to quantify the differences between species; for instance, studying functional homology across multiple areas simultaneously.

Another advantage of behaving rodent MRI is the relative ease of access to an overview of brain areas engaged in a task. For technical reasons, rodent studies typically focus on one or a few areas, which are decided based on previous studies. New studies tend to, on one hand, perpetuate the conditions under which an area was initially studied (e.g. within olfactory processing) and, on the other hand, overlook the importance of additional areas that have never been never probed. Our regression, decoding and functional coupling analyses reveal a range of task-related areas from early sensory to higher-order cognitive regions, highlighting the distributed nature of behaviourally-relevant information processing, echoing recent findings with electrophysiology (Steinmetz et al., 2019). Some of these areas were expected from the literature (e.g. body map in somatosensory cortex, primary visual areas in response to visual stimuli), but several others have either been largely unexplored or studied exclusively in a restricted set of conditions, consider for instance piriform cortex, which to our knowledge has never been studied in a non-olfactory task. Thus, behaving rodent fMRI offers an efficient and unbiased method to gain a global perspective over the brain areas engaged in a task (GLM analysis), their potential role (decoding analysis), and cross-region coupling (functional coupling analysis). This information can then be used to guide decisions about which regions to target, and to inform theories about how networks of areas work together to orchestrate behaviour.

There are a number of techniques available to investigate functional circuitry, each with its unique advantages and disadvantages. We chose functional magnetic resonance imaging for two main reasons: (1) because it allows observation of simultaneously recorded whole-brain activity during task performance and (2) because of its comparability with human brain studies where fMRI is most commonly used. Electrophysiology and calcium imaging offer high spatiotemporal resolution but are still far from achieving whole-brain simultaneous coverage (Lu et al., 2020; Sofroniew et al., 2016; Steinmetz et al., 2019). Histological marking techniques such as c-fos (Guenthner et al., 2013) allow whole-brain coverage at high spatial resolutions but are restricted to a single time point. Functional ultrasound (fUS) has recently emerged as a powerful combination of brain-wide coverage and high spatiotemporal resolution. However, acquisitions in awake rodents are still serial with different 2D slices being recorded at least 20 minutes apart (Macé et al., 2018). Importantly, none of these approaches can achieve whole-brain recordings in humans. By contrast, fMRI allows whole-brain simultaneous imaging across species from mouse to humans.

The relatively low temporal resolution of fMRI acquisitions is a limiting factor, however this can be partly mitigated by careful task design. The non-invasive nature of MRI means that there is virtually no upper limit on the number of sessions that can be acquired. Thus, one can start with low temporal resolution to gain a whole-brain overview, and upon observation, trade off spatial coverage for increased temporal resolution (e.g. by choosing a single slice in any orientation that can traverse several areas of interest) to reach temporal resolutions on the order of tens of milliseconds. Ultimately, one is limited by the sluggish nature of the haemodynamic process, in the order of seconds. However, BOLD onsets, measured using ultrafast BOLD sequences, are known to preserve the temporal order of neural events (Silva2002, Yu2012). Furthermore, with appropriate modeling of the haemodynamic response function, it will eventually be possible to estimate neural activity from BOLD and vice versa, as currently done for calcium imaging (Pachitariu et al., 2018; Theis et al., 2016; Vogelstein et al., 2010). Accurately modeling the haemodynamic response function is no easy task, but the use of tools available in rodents to record and manipulate with cell-specificity to investigate the principles of neurovascular coupling promise great and fast progress. Multiscale comparisons within the same animals have already been achieved in awake non-behaving rodents (Desjardins et al., 2019). By using a long-term cranial window that is MRI compatible, Desjardins et al. achieved optical access for microscopic imaging and optogenetic stimulation in the same animals that underwent BOLD fMRI. This approach will be instrumental in bridging BOLD fMRI signals to the underlying activity of neuronal circuits. Extending this approach to awake behaving conditions, such as the ones we present in the current study, will further allow comparison of these signals not only in response to sensory stimuli but in the context of complex cognitive behaviours translatable to humans.

We identified signal artefacts coupled to the contraction of muscles surrounding the skull during behaviour. To have a more accurate estimation of artefact timing, relative to the impoverished binary measure of infra-red beam break used for event detection, we included all the muscle surrounding the skull in our field of view during imaging. Given that muscle contractions induce field inhomogeneity and thus signal fluctuations, we were able to measure a proxy of muscle geometric changes. We then used the rich information contained in these muscle voxels to estimate both global and regional artefactual changes in brain fMRI signal. Given that BOLD signal related to neural activity is slow (in the order of seconds), we expect that removing brain signal correlated with the relatively fast muscle change, will preserve most of the true neural-driven signal. We focused on preprocessing strategies. It is possible that effective interventions earlier than preprocessing, such as data reconstruction or data acquisition, can be found. We used balanced steady steady free precession (bSSFP) due to its high spatiotemporal resolution and robustness to image distortions (Miller, 2012; Park et al., 2011; Zhou et al., 2012). In pilot studies, we tried two other acquisition strategies: Echo-planar imaging (EPI), which is more sensitive to BOLD at the cost of being more prone to artefacts, and Half-Fourier Acquisition Single-shot Turbo Spin Echo imaging (HASTE), less sensitive to BOLD but more robust to artefacts. In all, we saw artefactual amplitude changes coupled to licking. In EPI, they also led to mislocalization of the signal and thus apparent head motion. Importantly, realignment strategies were not sufficient to correct them. At the acquisition stage, it will be interesting to try more recent pulse sequences such as xSPEN (Zhang et al., 2017) shown to be extremely robust to field heterogeneities. Another possibility, assuming that the lick-induced changes reflect only fluctuations in the static magnetic field, would be to use dynamic shimming so that field changes are frequently compensated for.

The study of neural activity underlying rodent behaviour using fMRI is in its early days, but showing fast progress. Only three studies, to our knowledge, have performed fMRI imaging while simultaneously measuring behaviour (Han et al., 2019; Sakurai et al., 2020; Tabuchi et al., 2002). Tabuchi et al. measured fMRI responses on a section of the rat brain during drinking behaviour. However, they did not measure behaviour in the scanner, imaged only a coronal slice, and at minutes-scale temporal resolution, limiting the relevance of the approach to the study of brain-wide activity during task performance. More recently, Han et al. used fMRI to record brain-wide activity while mice performed a 2-odour olfactory go/no task (Han et al., 2019), while Sakurai et al. measured brain-wide activity during cued-water drinking. While these studies represented a significant advancement in the field, they have limitations that need to be tackled if this approach is to fulfil its potential as a valuable complementary tool for rodent behavioural neuroscience studies and a key technique in translational neuroscience. First, Han et al. and Sakurai et al. only record licking behaviour and deliver maximum 1 unimodal stimuli, offering limited ability to isolate sensory, cognitive, and motor processing. Instead, our setup allows for the use of multiple, multi-modal stimuli and three different behavioural readouts. Second, by employing designs with long task epochs (trial length ∼15s vs. 2s in previous studies), we were able to separate different task-related activity (e.g. correlates of preparatory and consummatory behaviour). Third, while Han et al. identify regions by averaging, and Sakurai et al. by using a univariate GLM, we establish the feasibility of a range of additional analysis strategies exploiting multi-voxel patterns and trial-by-trial variability, including multivoxel decoding and brain-wide correlational analysis. We thus sought to illustrate the breadth of information that can be obtained with this approach to tackle neuroscience questions in future studies. Finally, while Han et al. and Sakurai et al. only correct for potential artefacts arising from changes in head position, we dedicate substantial efforts to identify, characterise and correct artefacts arising from task-related jaw and body movements, which if left uncorrected can lead to false positives. It is possible that Han et al.’s and Sakurai et al.’s tasks and/or analyses are not sensitive enough to be affected by and pick up the artefact or that by focusing on activity that is different across groups (Sakurai et al., 2020), these artefacts were subtracted. Independently of the reason, it is paramount that these artefacts are thoroughly addressed, especially as we move from validation to exploration of richer and more complex tasks. A comprehensive method that will be widely applicable must therefore offer strategies to tackle not only artefacts arising from head motion but also other body movements. Our method exploits the rich information in muscle voxels to estimate and reduce both regional and global artefactual changes in brain signal induced by head muscle contractions. This represents an important step forward in ensuring valid and interpretable results in future studies, without requiring additional behavioural training or limiting task designs.

One of the main challenges in performing awake behaving fMRI is to minimise stress. Besides the possible stress arising from head fixation, MRI acquisitions lead to loud, variable noises and strong vibrations which can be sources of stress. Stress can not only lead to poor image quality due to head motion (King et al., 2005; Harris et al., 2015) but also impair learning and task performance (de Quervain et al., 1998; Diamond et al., 1994; Graham et al., 2010; Hölscher, 1999; Kaneto, 1997; Kim et al., 2001), being thus particularly crucial to minimise in awake behaving settings. Gradually exposing animals to MRI sounds has been shown to be effective at decreasing stress (Desai et al., 2001; Harris et al., 2015; Ferenczi et al., 2016; King et al., 2005). We combine this and additional steps to acclimate mice to the various potential stressors. Our procedure differs from previous behaving rodent fMRI methods (Tabuchi et al., 2002; Han et al., 2019; Sakurai et al., 2020) in several ways. First, it is longer and more gradual, aiming to minimize stress at all stages of training. Second, by training mice to voluntarily enter head-fixation, it avoids possible stress from manually fixation. Third, sounds are introduced only once animals are motivated and engaged in the task, and thus are less likely perturbed by them. Fourth, sound exposure is very gradual and lasts for at least 12 days as this was previously shown to be most effective (Harris et al., 2015). Finally, to habituate mice to vibrations, we also train them in the scanner. Despite our best efforts, it is possible that some residual stress remains. It will be important in future studies to complement these stress minimising procedures with external measures of stress (e.g. cortisol) as it has been done for sound exposure (Harris et al., 2015, King et al., 2015) and, when possible, directly compare MRI and non-MRI results in a specific region of interest.

The behavioural tasks we used were meant as a first step and were so far relatively unconstrained in terms of the cognitive processes engaged. It will be interesting, in future work, to extend this approach to richer and more constrained behavioural settings, where similar analyses can be used in conditions that only differ in a single quantifiable computational variable, to probe the underlying information representation in the neural systems. We extend the behavioural repertoire relative to previous awake behaving fMRI studies to include lever pressing in addition to licking as behavioural readouts. By requiring a movement different from reward consumption and separated in time from reward delivery, lever pressing offers a clear distinction both at the behavioural and neural level between the choice report and the reward consumption, facilitating the dissection of decision-making mechanisms. Furthermore, and contrary to licking, lever pressing is learnt, voluntary, and not reflexive, thus offering a more controlled and relevant readout for the study of choice behaviour. Importantly, the distinction between right and left lever press, although not exploited in our task, make our system compatible with more complex and controlled decision-making tasks, including two-alternative forced-choice, where psychometric models can be used to quantitatively relate stimulus features with behavioural choices.

In future olfactory studies, it will be important to complement the current approach with measurements of sniffing. Sniffing is known to be affected by olfactory stimuli (Laing, 1983; Sobel, 2000; Warren et al., 1994). For example in rats, the frequency and depth of sniffing depend on odour concentration (Youngentob et al., 1987). Furthermore, sniffing is known to be modulated by task variables such as reward expectations (Kepecs et al., 2007). This is important as fMRI signals will depend on blood oxygenation and thus respiration. Although the effect of changes in respiration would be expected to be global and not explain regional differences, it will be important in future studies to measure it simultaneously with fMRI data acquisition.

We strived to show how much can be obtained by having a rodent performing a task during fMRI. What we did not take advantage of was that, unlike in humans, in rodents we can acquire a high number of sessions and more easily access multiple subjects to improve signal to noise ratios when pursuing neuroscientific questions. This, in addition, offers the possibility of acquiring multiple data to use data-driven approaches to estimate HRFs more accurately. This was beyond the scope of our proof-of-principle study, but it will be a clear point of strength for future studies.

In conclusion, we describe protocols to train mice to perform reward-guided tasks in the scanner during whole-brain imaging with fMRI. We establish effective strategies to reduce inevitable artefacts arising from body movements and offer strong validation of these methods by mapping sensory, motor, and reward correlates in line with previous literature. We extend our approach to an operant task that separates consummatory and preparatory behaviour and offers higher design flexibility. Besides mapping task-related activity across the brain, we show how decoding analysis can be used to compare task-related activity across multiple areas, and how brain-wide correlational analysis can be employed to explore task-related inter-region coupling.

Our work illustrates how whole-brain fMRI data, combined with advanced analysis techniques, can be used to aid unbiased investigation of the brain-wide circuits mediating behaviour, paving the way to a whole-brain approach in rodent behavioural neuroscience. Combined with invasive manipulation techniques (i.e. optogenetics), it will enable tackling questions so far intractable even in rodents. How do cell-specific manipulations (e.g. optogenetic manipulations of neuromodulatory systems) influence brain-wide network activity in behaving mice? How do these global effects drive and are affected by behaviour? Finally, since fMRI is one of the cardinal techniques in human systems neuroscience, this approach offers a bridge from rodent to human neuro-behavioural studies, allowing direct investigation of similarities as well as differences between species.

## METHODS

### Animal subjects

Seven male adult C57BL/6 mice were used in this study. All procedures were carried out in accordance with the European Union Directive 86/609/EEC and approved by Direção Geral de Veterinária of Portugal. Animals (20-25 g) were group-housed prior to surgery and individually housed post-surgery. They were kept under a normal 12 hour light/dark cycle and tested at light phase. Mice had free access to food. Water availability was restricted to the behavioral sessions. Extra water was provided if needed to ensure that mice maintain no less than 75% of their original weight. Mice performed 1 session per day, 6 or 7 days a week. Five mice were used in the classical conditioning task. Two mice were used in the operant task.

### Stereotaxic surgery for head plate implantation

Animals were anaesthetised with isoflurane (4\% induction and 0.5-1% for maintenance) and placed in a stereotaxic frame (David Kopf Instruments, Tujunga, CA). Lidocaine (2%) was injected subcutaneously before incising the scalp. The skull was covered with a layer of Super Bond C&B (Morita, Kyoto, Japan) to help stabilize the implant: 3 plastic nuts (M2, PKN2, Solid Spot, Santa Clara, CA, USA). The implant was cemented to the skull using dental acrylic (Pi-Ku-Plast HP 36, Bredent, Senden, Germany). Mice were monitored until recovery from the surgery and returned to their home cage where they were housed individually. Gentamicin (48760, Sigma-Aldrich, St. Louis, MO, USA) was topically applied around the implant. Water deprivation and behavioral training started at least one weeks after surgery.

### MRI compatible behavioural setup

Progress in whole-brain imaging in behaving rodents has been hampered by the hardware constraints of MRI scanners. The strong magnetic field and the restricted space in the scanner means that the solutions typically used for rodent fication (e.g. metal head plates) and behaviour quantification (.e.g large metallic components) are problematic. To overcome these obstacles, we combined fibre optics, pressure sensing, and 3D printed plastic parts into an MRI compatible setup that delivers multiple sensory cues (i.e. olfactory and visual), water rewards and records multiple behavioural measures (licking, right lever press, left lever press).

Head-fixation was achieved by combining a plastic (PEEK) nuts (M2, PKN2, Solid Spot, Santa Clara, CA, USA), chronically implanted on the mouse’s skull, a custom-made 3D printed fixation platform (designed and printed in-house) secured to the MRI bed and three plastic screws (M2, PEEK/PH M2-4, Solid Spot, Santa Clara, CA, USA) that secured that connected the implanted nuts and the platform.

Lever pressing was detected using two infra-red beam break systems delivered and sensed via long-range optic fibres. Each beam-break system was composed of two custom designed fibers (“U-fibre-light-detector”, Doric lenses, Quebec, Canada), a custom IR emitter and control unit (“Custom Control/Detection Electronics for U-Bracket”, Doric lenses, Quebec, Canada), and an amplified photodiode for detection (APD_FC, Doric lenses, Quebec, Canada). In the mock setup we used infra-red emitters (SEP8736-003, Honeywell, Bracknell, UK) and phototransistors (480-1958-ND, Honeywell, Bracknell, UK).

Licking was detected using a pressure sensor that detected lick tube (EW-06407-41, Cole-Parmer, Vernon Hills, IL, USA) vibration via an air pad (Respiration pillow sensor, SA Instruments Inc., USA), positioned under the tube holder, and connected to a monitoring system (Model 1030 Monitoring Gating System, SA Instruments, USA). The tube holder was 3D-printed in house (Hardware Platform, Champalimaud Research, Portugal). In the mock setup, we used a non-MRI compatible infra-red beam break detector (GP1A57HRJ00F, Sharp, Uxbridge, UK).

Odours (amyl acetate, 1-hexanol, ethyl butyrate, eugenol, diluted in mineral oil in a 1:10 ratio) were delivered using a custom-built olfactometer. Diluted liquid odours and a blank control odour (pure mineral oil) were loaded onto disposable syringe filters (20 μl, Whatman) that were then inserted into a PEEK manifold. The flow of air through each of the filters was controlled independently using a three-way solenoid valve (360K031, NResearch, NJ, USA). At any given moment, only one of the valves was open, passing either odourised or blank air streams at a flow rate of 100 ml/min through the filter and into the manifold. This stream was mixed within the manifold with a second, “carrier” stream of clear air (flowing at 900ml/min; both streams controlled independently with flow meters, regularly calibrated using a mass flow meter (Model GFM17A-VAL6-*O, Aalborg) from which it was delivered through Teflon tubes (EW-06407-41, Cole-Parmer, Vernon Hills, IL, USA) to the animal’s nose. Photoionization detector measurements showed that odour was sensed in less than 250 ms delay from valve opening.

Visual stimuli were delivered via optic fibres connected to fibre-coupled LED sources. Visual stimuli (blue or yellow light) were delivered via long-range optical fibres (200 μm, 0.22 NA, Doric lenses, Quebec, Canada) connected to fibre-coupled LED (470nm, CLED, Doric lenses, Quebec, Canada), which was in turn, connected to a LED driver (LEDD1B, Thorlabs, Newton, NJ). The tip of the fibre delivering yellow light was positioned close to the mouse’s right eye, the tip of the fibre delivering blue light was positioned close to the mouse’s left eye.

Water was delivered using a distant water valve (LHDA1231215H, The Lee company, Westbrook, CT, USA). All systems connected to an Arduino-based board that was controlled by a dedicated computer using Arduino software, Python and Bonsai.

All the electronic and metal components were positioned at a safe distance from the scanner and connected to the setup through long-range connections.

This system was designed for use with a cryogenic surface array coil at 9.4T, but is in principle compatible with other coils. In pilot studies, we used a similar system for non-cryogenic array surface coil using a previous version of the head-fixation system and a beam-break lick detection (Costum U-bracket, Doric lenses, Quebec, Canada) using an IR led source and long-range optical fibre.

### Classical conditioning task

Each trial started with the presentation of one of four odors for 2 s, randomly selected. Following a 3 s fixed delay (trace period), the corresponding outcome was available. For two odours the outcome was a water reward (∼4 microliters) and for the other two it was nothing, counterbalanced across mice. Outcome delivery was followed by a variable inter-trial interval (6:9s, drawn from an uniform distribution). Mice were first trained only with the rewarded odours with an odor period of 1s, trace delay of 0s and ITI period of 2-3s. Neutral odours were introduced once mice showed anticipatory licking (licking during the trace period). The odour period, trace period and ITIs were increased gradually during training until the final values: 2s odour presentation, 3s trace and 6-9 ITI.

### Operant conditioning task

A trial started with a yellow cue signaling the waiting period (2-4s, increased over sessions) during which presses were not rewarded. The first press (either right or left) after the waiting period elapsed, was defined as a “valid press’’ and was rewarded after a delay (0.5 - 1.5s, increased across sessions). A blue light provided immediate positive feedback after a valid press and stayed on until reward delivery, serving also as a bridging cue. Reward delivery was followed by a short intertrial interval (0.5s). During the initial stages of training every single press (left or right) was reinforced with a water reward. This produced reliable pressing but meant that mice would become satiated very quickly and that licks were tightly coupled with presses. To slow down the behavior and start decoupling presses from reward, and thus licking, we inserted the cued waiting period (0.5-4s) during which presses were not rewarded (effectively an inter-trial interval). The delay to the reward was also increased over sessions. To promote variability between individual presses and between presses and licking, we avoided overtraining and even within experimental days, we increased the length of all task delays over sessions.

### Training procedure

A major challenge for doing simultaneous fMRI and mouse behaviour is to train mice to be comfortable enough in the scanner to perform successfully behavioral tasks during imaging and without disrupting image quality with head motion. To achieve this, we used a mock setup replicating the one in the scanner and developed a sequential training procedure that exposed mice gradually to the various components (i.e. noise, vibrations) that would be present in the functional imaging sessions.

After surgery for head holder implantation and recovery (1 week), water deprivation began. Following 2-3 days of handling in the homecage (Guo, Hires, et al., 2014), mice were trained to voluntarily enter the head-fixation apparatus. The motivation for this choice is two-fold: firstly, to minimise any fixation-related stress, and secondly, to avoid the use of anaesthetics, which are known to dampen fMRI signals and take tens of minutes (20-30 mins) to wash out even after a brief initial use.

In the first day, mice were allowed to freely explore the mock setup without the head-fixation screws. At the beginning of the session, the water spout was easily accessible without requiring mice to insert the head all the way through the platform. After some successful approaches, the spout was gradually moved further away, until reaching its final position.

The following day, one long screw was attached to the most anterior head nut so that the licking spout was only accessible if the screw entered the platform’s slit. Mice were allowed to freely remove themselves from the fixation platform and re-enter. Each entry was reinforced with a water reward. Maintaining the head in the platform led to additional rewards (up to 10 rewards, 1/second). Typically mice started by doing very short entries and progressively stayed longer. The procedure was repeated for another day with a shorter screw, making slit-entries more difficult, and the number of rewards per entry were increased to a maximum of 20, to further reinforcing staying in the platform. In the final stage, mice entered the holder and the head screw was tightened for increasingly longer periods. If at any point mice exhibit stress signs such as struggling or vocalisations, mice are immediately released and the session was terminated. This phase lasted 8-10s days.

Next, mice underwent task specific training (8 days for the classical conditioning task, at least 2 weeks for the lever press task) and, once proficient, were exposed gradually to increasingly loud MRI sounds recorded from the scanner. The volume of the sound was increased gradually from session to session, in at least 5 steps. Mice were exposed to the maximum loudness for at least 7 more days. We monitored continuously the animal being especially attentive to any struggling movements, vocalisations or interruptions of licking. Typically, in the first 1-2 days of sound exposure, mice interrupt licking at sound onset and after a few seconds resume it.

After undergoing sound acclimation, mice were taken to the MRI room, where they were first trained outside the scanner to allow habituation to the new environment and then inside the bore (2 days) with the real scanner sounds and vibrations. Each experimental session started with a few “dummy trials” to keep mice engaged as the setup was inserted into the scanner bore and the imaging adjustments were performed. After adjustments, functional images were acquired. Reference anatomical images were acquired at the end of the session. The entire procedure lasted 30-40 days for the classical conditioning task, longer for the lever press task.

### Magnetic resonance imaging

#### Small animal scanner and radiofrequency coil

All MRI experiments performed on a 9.4 T Bruker BioSpec MRI scanner (Bruker, Karlsruhe, Germany) equipped with an AVANCE III HD console including a gradient unit capable of producing pulsed field gradients of up to 660 mT/m isotropically with a 120 us rise time. A 86 mm quadrature resonator was used for transmission, and a 4-element cryoprobe (Bruker, Fallanden, Switzerland) was used for reception. The software running on this scanner was ParaVision 6.0.1.

#### Subject preparation

On the day of the experiment, the mouse was taken from the home cage on the palm of the hand of the experiment. The plastic screws attached to the chronically implanted nuts were carefully loosened. The mouse was then placed in the MRI behavioural bed, in the scanner room, and allowed to freely explore the setup. A few drops of water were placed forming a path to the head-fixation platform and the water spout. Once the mouse entered the head-fixation platform through the slit, the experimenter tightened the screws, and the task “dummy trials” began. Finally, the bed was inserted into the scanner and the imaging adjustments began. After adjustments, functional images were acquired. Reference anatomical images were acquired at the end of the session, when mice were satiated.

#### Scanner synchronised behavioural monitoring

An arduino MEGA260 connected to the LED source and the laser, and receiving triggering from the MRI scanner, was used to generate pulses of LED light or laser light, respectively.

### MRI data acquisition

#### Adjustments

The head-fixation platform ensured reproducible, optimal positioning in the scanner. Once the animal was positioned in the scanner, localizer images were acquired to assess the quality of the position. A B0 field map was obtained from phase images acquired using a 3D-dual-gradient-echo pulse sequence. The correction of B0 field inhomogeneities was automatically performed using a MAPSHIM routine in a cuboid volume comprising ∼525 mm^3^ located in the brain and centered on the middle slice of the fMRI acquisitions.

#### Anatomical scans

In the lever press classical conditioning task, anatomical images were acquired using a T2-weighted Turbo Rapid Acquisition with Relaxation Enhancement (RARE) sequence (TE_eff_/TE/TR = 32/16/4500 ms, RARE factor = 6, Number of averages = 1, FOV 15 x 18 mm^2^, Matrix size = 100 x 120, in-plane resolution = 0.15 mm^2^, slice thickness = 0.8 mm, Number of slices = 10). No anatomical images were acquired in the classical conditioning task.

#### Functional scans

Functional scans were acquired using a fast-imaging with steady-state precession (FISP), also known as balanced steady-state free precession (bSSFP) sequence. FISP was chosen due to its high spatiotemporal resolution and robustness against image distortions (Miller, 2012; Park et al., 2011; Zhou et al., 2012). In the classical conditioning task we used the following parameters: TR/TE = 2.3/1.6 ms, 1 shot, Flip angle 30°, FOV 15 x 13, Matrix size 76 x 66, in-plane resolution 0.2 x 0.2 mm^2^, number of slices = 9 coronal slices with a 0.15 mm gap, slice thickness 0.75 mm. In the lever press operant conditioning task, we used similar parameters adjusted for differences in resolutions: TR/TE = 2.8/1.4 ms, 1 shot, Flip angle 30°, FOV 15 × 18, Matrix size 50 x 60, in-plane resolution 0.3 x 0.3mm^2^, number of slices = 10 coronal slices with a 0.2 mm gap, slice thickness 0.8 mm.

### Behavioural data analysis

#### Linear temporal drift

We observed a time drift between the Arduino and the MRI clocks. The time drift appears evident when considering the time course of the voxels belonging to the part of the volume occupied by muscles, with respect to the time series of events detected by the Arduino board (e.g. presses, licks).

To avoid overfitting and hence bias the subsequent analyses, we assumed the drift to be linear in time. A straightforward algorithm to re-synchronise the signals is the following:

1. *E*(*t*) by multiplying the indices of *E*(*t*) by *m* .identify the onsets of the bouts of muscle activity *M*(*t*) and detected events *E*(*t*), respectively. Let 𝒪(*M*) and 𝒪(*E*) be the collections of detected onsets.
2. Find the one-to-one function *φ* : 𝒪(*M*) → 𝒪(*E*) such that the distance |*μ* − *φ*(*μ*)| is minimal for every *μ* ∈ 𝒪(*M*).
3. Regress *m* (*y* = *mx*) by considering the point cloud {(𝒪(*M*), *φ*(𝒪(*M*)))}.
4. Warp *E*(*t*) by multiplying the indices of *E*(*t*) by *m*.

We use this strategy to support the validity of the linearity assumption, as shown in **Figure S5**. However peak detection can suffer from the noisy nature of our signals, requiring heavy, arbitrary preprocessing. For this reason, we chose such that it maximises the correlation at time 0 between the mean in time of the absolute value of the muscle activity and *E*(*t,m*). To do this, we consider 100000 evenly spaced values for *m* in the closed interval [0.98, 1.01]. Thus, if *m* = 1 maximises the correlation, no drift is detected and the optimal warped event time series would correspond to the original one. **Figure S6** shows the output of the drift-correction algorithm. The correlation-maximisation procedure is detailed in the caption.

#### Conversion from time to frames

Behavioural parameters are downsampled to match the frame rate of acquisition of the scanner. This is done by associating to the i-th component of the vector the number of occurrences of the behaviour in the timespan necessary to acquire the frame. Clearly, behaviours occurring at a maximal rate equal or inferior to the one of slice acquisition will be associated to boolean vectors. Finally, the downsampled behavioural vectors are divided into task and nuisance regressors.

#### Classical conditioning task

Anticipatory licking was defined as any licking that occurred after mid-odour presentation until just before outcome presentation. Average lick rate traces were computed by smoothing the lick counts using convolution with a Gaussian filter of 50 ms standard deviation. Mean lick rates were computed by taking the number of licks within a defined window (750:0 s for anticipatory licking and 100:850 ms for consummatory licking, all times relative to outcome delivery), divided by the duration of that window. To correct for differences in overall lick rate between scanner and non-scanner sessions due to the use of different detectors (beam break electrical detector in the mock setup and motion sensor in the scanner), the mean anticipatory or consummatory lick rates were divided by the total lick rate in the session (total number of licks divided by the session duration). Mean lick rates were either computed per odour or pooling odour with the same reward contingency, as indicated in the text.

#### Operant conditioning task

Valid presses were defined as the first presses after the waiting period. Invalid presses were defined as any press during the waiting period. Latency to press was defined as the time between the end of the waiting time and the first press. Short-latency presses were defined as valid presses with latencies below the 50th percentile of all press latencies for that animal. Long-latency presses were defined as longer than the 50th percentile.

### MRI data analysis

We processed the data following standard fMRI data processing. We tackled the correction of artefacts, arising from changes in muscle configuration, using a custom strategy. Our main approach to the correction of these artefacts was task-agnostic and based on LASSO regression, as described below. In the remainder of this section, we describe each of the processes involved in the analysis pipeline of both images and behavioural data. The algorithm flow is described in **Figure S7**.

#### Standard preprocessing

Functional images were preprocessed using custom-written scripts in MATLAB and Python, and scripts from SPM12 (Wellcome Trust Centre for Neuroimaging, London, UK, http://www.fil.ion.ucl.ac.uk/spm/).

First, functional and anatomical images were converted from Bruker format to NIFTI format. Images were then corrected for slice-timing differences using sinc interpolation with the SPM12 toolbox. This is to correct for the fact that different 2D slices composing a 3D volume are acquired at slightly different timings. Next, images were realigned using rigid-body transformation to correct for motion-related changes in brain position. For population analysis, images were co-registered to the subject’s own T2 anatomical image and normalized to a common space (one of the animals). Finally, to correct for scanner low-frequency drift and for absolute signal differences arising from coil placement (signal decays with distance to the coil), the mean and linear trend terms were removed from each voxel’s time series using voxel-wise linear regression.

#### Lick artefact correction

We found signal artefacts coupled to lick events and changes in muscle tissue. We aimed to use the information contained in the voxels in the muscle to predict the artefact in the brain, as muscle changes correlated better with artefact than the event detection vector. This is because muscle contractions can start before and extend slightly after an event detection, measured as a beam break or spout movement (e.g. opening of the mouth before the tongue protrusion that touches the spout). We chose least absolute shrinkage and selection operator (LASSO) (Tibshirani, 1996), as we had a large number of predictor (muscle) voxels. LASSO is a regularisation method that by pushing to zero the coefficients of voxels that are least predictive, performs covariate selection, using only a subset of voxels in the final model. **Figure S9** shows the contribution of each muscle voxel to the predicted intensity of the brain’s voxels for two example sessions. Note the small number of contributing voxels. In the following sections we describe the basic definitions needed to introduce LASSO regression, then we provide details concerning the use of LASSO regression in our artefact removal algorithm.

##### LASSO regression

Let *D* be a dataset consisting of {(x_1_, *y*_1_), …, (X*_N_*, *y_N_*,)} observation and expected value pairs. For every *i* ∈ {1, …, *N*}, we have that x*_i_* = (x_1_, …, *x_m_*) ∈ ℝ*^m^* and *y_i_* ∈ ℝ. LASSO regression minimises the objective function:

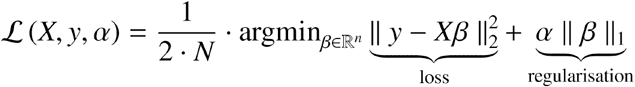

When minimising ℒ, we fit a linear model of *y* on *X*, and regularise the regression coefficients by using their *L*_1_ norm. The strength of the regularisation is proportional to the parameter *α*. Let, **v** = (*v*_1,_ …, *v_m_*) ∈ ℝ*^m^*, the *L*_1_ norm of **v** is defined as:

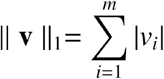

*L*_1_ regularisation favours coefficients to be zero as shown in **Figure S8**. This feature, when compared to other kinds of regularisations, makes LASSO an ideal strategy for feature selection.

It is also possible to generalise this technique to multidimensional expected values, i.e. when *y* ∈ ℝ*^l^* with l > 1. In this case the loss function can be rewritten as:

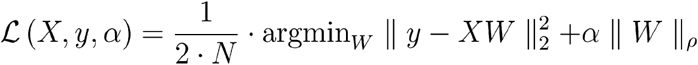

where *W* ∈ Mat (ℝ, (*N*, *l*)), ||·||_2_, is in this case the Frobenius norm for matrices and ||·||_ρ_ is computed as the sum of the 1-norm of each row.

##### LASSO regression for artefact correction

Muscle contractions occurring during behaviour introduced artefacts in the recording. Our main approach to the correction of these types of artefact was task-agnostic and based on LASSO regression. After the standard MRI preprocessing procedures (slice-timing correction, motion correction and drift removal), we use the following steps. First, we manually generate for each slice *s_i_* the masks *M_i_* and *B_i_* associated to muscle and brain voxels respectively. Each *M_i_* is chosen conservatively, in order not to include any voxel entirely or partially belonging to the brain area. Let *M_ij_* and *B_ij_* be the masked images obtained by applying the muscle and brain mask to the slice at frame *j*. All the masked images are standardised (centred and divided by their standard deviation). Then, the collection of *M_ij_* is used as a set of observations, and *B_ij_* as target values in the computation of the multivariate LASSO regression ℛ.

Essentially, we compute at each frame *j* how each voxel in *M_ij_* contributes in explaining the activity recorded in *B_ij_*. The activity explained by *B_ij_* is then subtracted from *B_ij_*. In symbols, a corrected image is computed as *C_ij_* = *B_ij_* − ℛ(*M_ij_*). Finally, we denormalise the corrected images by adding the mean of *B_i_* along the entire session.

#### General linear model

To reduce thermal noise and compensate for imperfections in co-registration, images where smoothed in space using a gaussian kernel with a full-width half maximum (FWHM) of twice the voxel size (0.3 mm for classical conditioning task and 0.6 mm for the operant task) before GLM fitting (Friston et al., 2007). For the statistical t-maps, GLMs were fitted either per subjects or pooling various subjects. Given that our N was small we used only fixed-effects analysis. We convolve behavioural events (CS+,CS-,US+,US- in the classical conditioning task; or right, left and licks in the lever press operant task) with the canonical haemodynamic response function (Glover, 1999) (a difference of two gammas). This is likely suboptimal for rodent analysis, however there is still little data on event-related fMRI in rodents to adequately estimate the HRF. For this reason, we conservatively used the canonical HRF, well characterised in humans. Future studies using data-driven approaches to estimate HRF will be important to increase the sensitivity of these analyses. To reduce false-positive activation generated by residual motion, we included the 6 motion correction parameters (x,y,z coordinates and pitch, raw, yaw rotations) generated in the realignment preprocessing step. In addition, we extracted the intensity time course of representative voxels in the ventricles (cerebrospinal fluid, CSF), thought to reflect physiological and non-physiological noise (e.g. vascular pulsation, respiration). The first 10 volumes were skipped because the steady-state in the FISP was only reached approximately at volume 5. For all GLM analysis, the appropriate SPM contrasts were built in order to obtain maps for each task-related response. In the classical condition, we contrasted trial types (e.g. rewarded versus neutral trials) [1 -1], in the operant task, we contrasted with the baseline [1 0]. The resulting statistical t-maps were corrected for multiple comparison using family-wise error rate (FWE) corrections for p < 0.05 and a minimum cluster size of 8 voxels (Friston et al., 2007).

#### Event-locked averaged maps

We aligned images to the specific event class (e.g. odour A, right lever press), baseline subtracted them with the mean image in the interval -2:-1s relative to event onset and averaged over events to generate an averaged map over time (Figure 6B), or averaged over timepoints (0:10s from odour onset) to generate an overall response map (Figure 4C). The main advantage of this approach is that it does not need convolution with an haemodynamic response function (HRF), thus allowing for an unbiased —albeit clearly less expressive— analysis. For the lever press maps aligned on right press for mouse #1, the event selection was further constrained to exclude any right presses that were either preceded or superseded by a left press in a window of -10 to 10 frames.

#### ROI analyses

A C57BLJ mouse atlas from the Allen Brain Atlas (ABA) was manually spatially warped to match the mouse reference images. ROIs were defined manually according to the atlas for the ROI averaged time courses and automatically generated for the decoding and functional connectivity analysis.

For the averaged time course analysis, we extracted voxels from the selected ROI (already detrended during preprocessing). Time courses were averaged over voxels, aligned on the specific event, baseline corrected (-2:-1s from event onset) and averaged over instances of that event class. In the lever press averaged time course, averaging over the two mice tested, the additional exclusion of right events based on left presses was not used as this decreased significantly the number of trials in mouse #2.

#### Decoding

We decoded both task-specific and behavioural events by using standard machine learning techniques and cross validating with stratified 5-fold. For odour identity decoding we used Multinomial Logistic Regression, as we had a small dataset per animal (1 session per animal, 10-20 instances per odour per animal). For the lever operant task, we used Support Vector Machine algorithms, since we had a larger dataset per animal (7 sessions). For all decoding analysis non-smoothed images were used. F1-score is defined as 2 * (precision * recall)/(precision + recall).

In the classical conditioning task, decoding was performed in the olfactory bulb for each animal, each odour and for each time point in a window from 0 to 10 frames from odour onset. To account for possible differences in response timing, we took the maximum F1-score across those delays. Those values were then averaged across mice. In the operant task we performed decoding for multiple ROIs for each animal, pooling data from all sessions. We took the maximum within the window used for each type of decoding (1:5 after press for valid vs. invalid press decoding and -3:3 for the decoding of short- vs. long-latency presses). We ranked regions according to their F1-score for each animal. We took the top (bottom) 15 regions with highest (lowest) F1-score for each animal. Among those sets, we found the regions that were common across both animals. For example, if region X ranked 2nd for one animal and 5th for other, it was selected. If region Y ranked 12th for one animal and 20th for another it was not included. We then plotted the 5 highest and 5 lowest from this common pool with an individual bar per animal for each of them. In the following sections we describe the definition of the classification algorithms we used, provide their geometrical intuition, and detail their parameterisation.

##### Decoding preprocessing

The training dataset is generated from drift and artefact corrected images and events, obtained via the procedures described in previous sections. First, images associated with events to be decoded are selected, allowing for a time shift to account for the slow nature of the BOLD response. Let 𝒟 = {(*i*_1,1,_ *e*_1,1_), …, (*i*_1,*n*1,_ *e*_1,*n*1_), …, (*i_k_*_,1_*e_k_*_,1_), …, (*i_k_*_,*nk*,_ *e_k_*_,*nk*_) be the images, event pairs collected for each considered event, and *n*\* = min ({*n*_1_, …, *n_k_*}). We consider the balanced dataset D = {(*i*_1,1,_ *e*_1,1_), …, (*i*_1,*n*\*,_ *e*_1,*n*\*_), …, (*i_k_*_,1_*e_k_*_,1_), …, (*i_k_*_,*n\**,_ *e_k_*_,*n\**_)}. Finally, we generate the 5-folds used for cross validation.

##### Stratified K-fold cross-validation

We split *D* in train and test set by partitioning it in *K* sets *D* = {*S*_1_, …, *S_k_*} of equal cardinality. Classifiers are trained on all the *K* possible collections *D_i_* = {*S*_1_, …, *S_i_*_−1_, *S_i_*_+1,_ …, *S_k_*} obtained by removing the *i* th subset of *D* and validated on *S_i_*. Statistics on these *K* training and validation procedures are used to assess the robustness of the decoding.

##### Multinomial Logistic Regression

We provide a geometrical intuition on logistic regression and the basic definition of multinomial logistic regression. We refer the reader to (Böhning, 1992) for more details.

###### Logistic regression

Let us consider a binary classification problem, i.e. a set of *n* labelled samples *T* = {(*x*_1,_ *y*_1_), …, (*x_n_*, *y_n_*)}, where (*x*_i,_, *y*_i_) ∈ ℝ*^m^* × {−1, 1}. The logistic function (sigmoid) is defined as 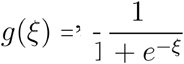, where 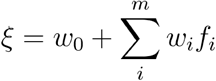, where *f_i_* are features associated to each sample, so that *x_k_* = (*f*_1_, …, *f_m_*) for every *a* = 0, *b* = 481, *f*(*x*) = *A* (*x*). Geometrically, the algorithm works by adding a new dimension for the dependent variable and fits a logistic function to {(*x_i_*, *y_i_*)}*_i_*_∈*I*_ so to separate samples for 0 ≤ *g* (*ξ*\*) ≤ 1. See **Figure S10B**. Given a new sample *x*, it can be classified simply by evaluating *g* (*ξ*\*(*x*)) and assigning the sample to class if *g* (*ξ*\*(*x*)) ≥ 0.5 and to class 0 otherwise.

###### Multinomial logistic regression

We utilised multinomial logistic regression for decoding images associated to more than two events. The multi-class equivalent of the logistic function is the softmax function

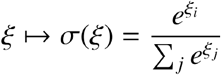

The softmax function outputs a probability distribution, thus the predicted events can be chosen by simply considering argmax σ(*ξ*).

##### Support Vector Machine

In a binary classification task, Support Vector Machines (SVMs) find the maximum margin hyperplane that separates the samples belonging to the two classes (Suykens & Vandewalle, 1999). See **Figure S10C** for an intuition. Let *T* be the labelled dataset described in the previous section, the margin hyperplane can be found via minimisation of the function

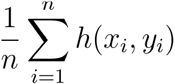

where *h* : (*x_i_*, *y_i_*) ↦ max(0, 1 − *y_i_*(*w* · *x_i_* – *w*_0_)) is the hinge loss function, where · is the dot product between the weights and the sample’s features.

The kernel trick (Boser et al., 1992) generalises the SVMs framework to nonlinear classification. This strategy consists in substituting dot products in *h* with a nonlinear kernel. We used a homogeneous polynomial kernel of degree 3, and kernel coefficient 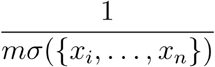, where *C*_3_*H*_8_(*g* + 5*O*_2_(*g*) → 3*CO*_2_(*g*) + 4*H*_2_*O* is the number of features and is the standard deviation.

#### Functional coupling analysis

Non-smoothed images were used for these analyses. For seed-based correlations, we computed for each session the Pearson correlation between a time series of an averaged seed ROI and each brain voxel. For voxel or ROI-wise correlations, we computed the Pearson correlation coefficient between the time series of a voxel with every other voxel, or between the average time course within an ROI with the averaged time course of every other ROI. We then averaged the resulting coefficients over sessions (N = 7). If each variable has N scalar observations, then the Pearson correlation coefficient is defined as:

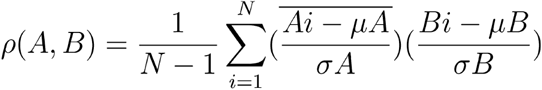

where *μA* and *σA* are the mean and standard deviation of A, respectively, and *μV* and σ*B* are the mean and standard deviation of B.

The chord diagrams shown in **Figure 8C-D** are available in a web-based interactive form at https://madalena_fonseca.gitlab.io/coupling_visualisation/.

## AUTHOR CONTRIBUTIONS

M.S.F., N.S., and Z.F.M. designed the experiments. M.S.F. developed the behavioural setup, with assistance from the Champalimaud Hardware Platform, and training procedures. M.S.F. conducted all the experiments. M.S.F, N.S., Z.F.M., and M.B. designed the analysis. M.S.F. conducted all analyses except the lasso regression and decoding, with input from M.B. M.B. performed the lasso regression and decoding analyses with input from M.S.F. M.S.F. wrote the manuscript with input from Z.F.M, N.S. and M.B.

## ACKNOWLEDGEMENTS

We thank the Masayoshi Murakami, Eran Lottem, Dario Sarra, Cindy Poo, Bass Atallah, Cristina Chavarrias, Daniel Nunes and Teresa Serradas-Duarte for daily discussion; Dario Sarra, Sofia Soares, Sam Walker and Teresa Serradas-Duarte for input on an earlier version of the manuscript; the Champalimaud’s Hardware platform, in particular Paulo Carriço for help developing and building the setup; Sebastião Van Uden and João Luis for help with initial 3D prototypes; Bruno Cruz and Pedro Garcia da Silva for technical assistance; João Cruz, Vesna Petojevic, Catarina Pimentel and Margarida Nunes for logistic support; the Champalimaud’s vivarium staff for experimental support. This work was supported by Fundação para a Ciência e Tecnologia (SFRH/BD/52446/2013, M.S.F.), European Research Council Advanced Investigator Grant (ERC-2016-AdG-671251, Z.F.M.) and Starting Grant (679058, N.S.) and Champalimaud Foundation (Z.F.M. and N.S.).

## SUPPLEMENTAL INFORMATION

**Figure S1.**
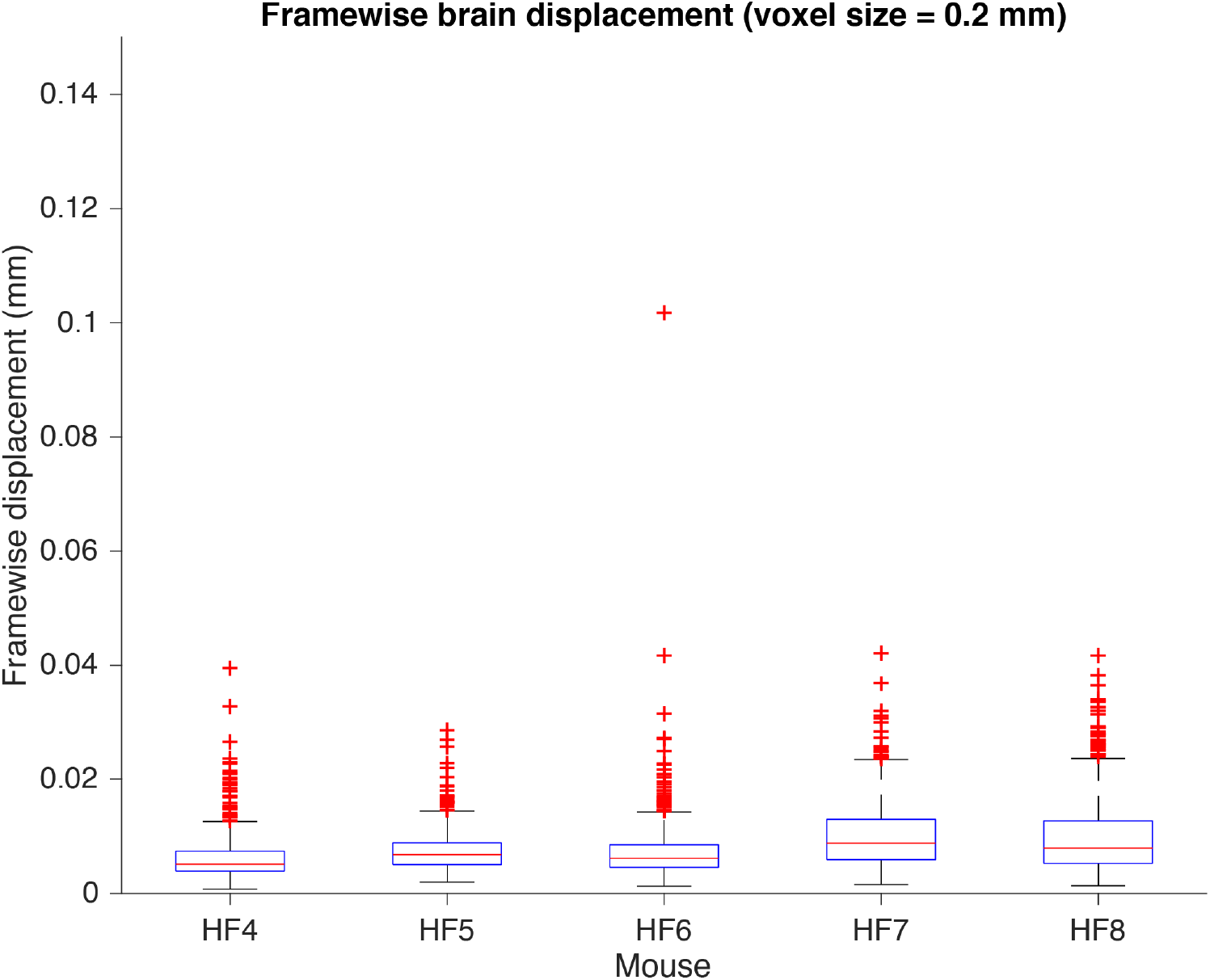
Related to Figure 3 - Framewise brain displacement. Boxplots with framewise displacement for all imaging frames (600-900 frames) for each mouse prior to any artefact correction. Boxplots show minimum (lower black line), first quartile (lower blue line), median (red line), third quartile (upper blue line), maximum (upper black line) and outliers (red cross). Frame-wide displacement was computed using 3 translations and 3 rotations (assuming a mouse brain radius of 5mm) parameters using SPM’s realignment function. We masked the muscle voxels leaving only brain-voxels in the input images to avoid muscle movement affecting the realignment. It is likely that these are still overestimations of brain movement as realignment strategies are intensity-based and thus can be influenced by the amplitude changes caused by lick artefacts.

**Figure S2.**
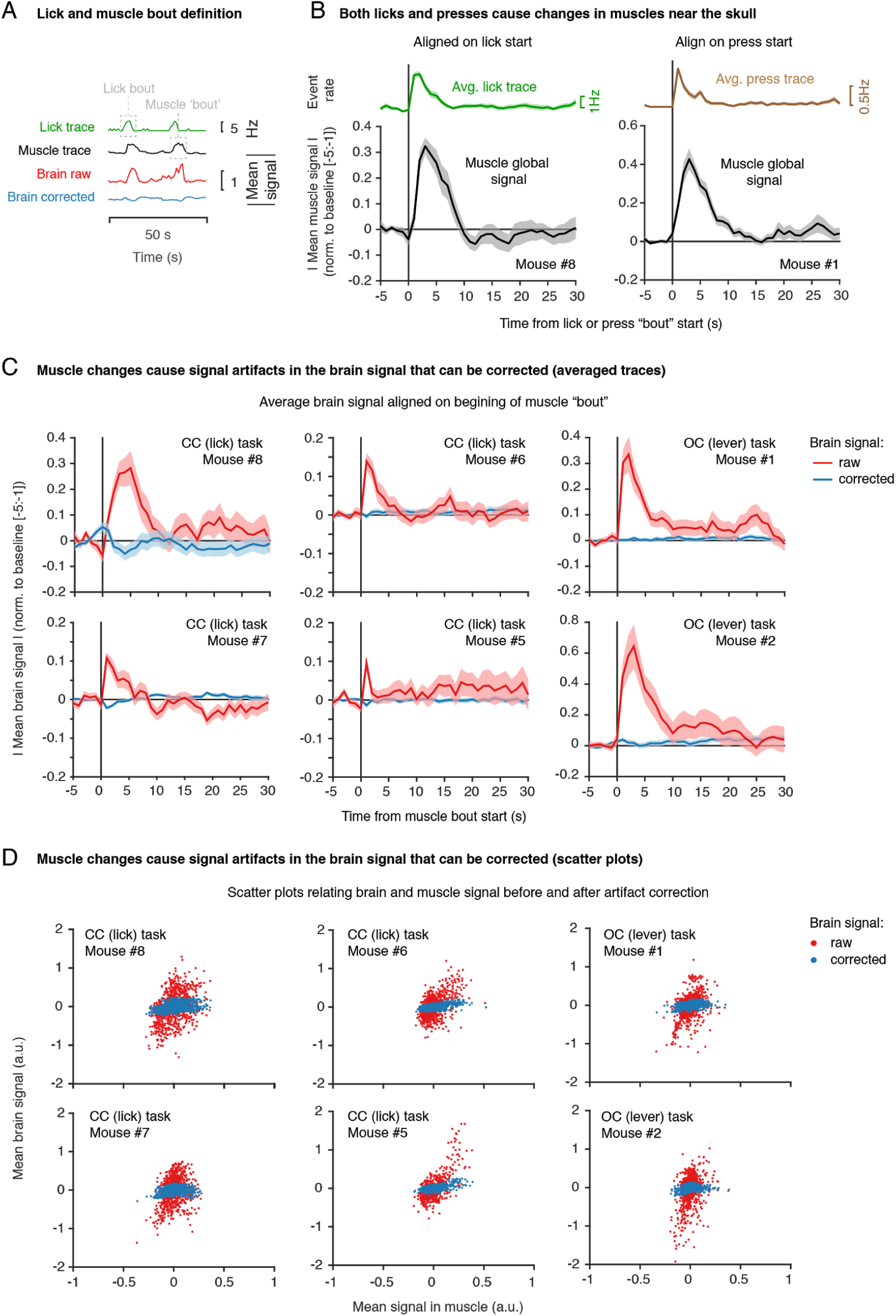
Related to Figure 3 - artefact correction in the classical conditioning (CC) and operant conditioning (OC) task. **(A)** Lick and muscle bout definition. Shown are four traces over time (for 50 s of the example session shown in Figure 3) for lick rate, muscle global signal, brain global signal before artefact corrections (raw) and brain global signal after correction (corrected). Grey boxes illustrate in the lick and muscle trace what is defined as a bout (consecutive lick events or muscle movements). Muscle global signal is an average over all voxels outside the brain but within our field of view (as shown in Figure 3), including tongue, jaw muscles and temporal muscles. For information on the artefact (LASSO) regression see **Methods. (B)** Averaged muscle global signal (signal averaged across voxel and across bouts, in black), aligned on either the start of a lick bout (left, lick rate averaged trace shown above in green) or lever press (right, lever press averaged trace shown above in brown). Note that both licks and presses cause similar changes in muscle global signal. Shown is one representative session. **(C)** Averaged global signal in the brain aligned on the beginning of a muscle bout for individual mice (the 4 mice considered for task-related fMRI analysis in the CC task and 2 mice in the OC task) in a single session. Where multiple session data were available, one example session was selected. In red is the signal before correction (raw), in blue is signal after correction (corrected). Muscle changes correlate with artefacts in brain signal (red, raw) that can be corrected (blue, corrected). **(D)** Scatter plots for the same individual mice relating brain and muscle signal before artefact correction (red, raw) and after correction (blue, corrected) for a single session. artefact correction diminishes the correlation between brain and muscle signal.

**Figure S3.**
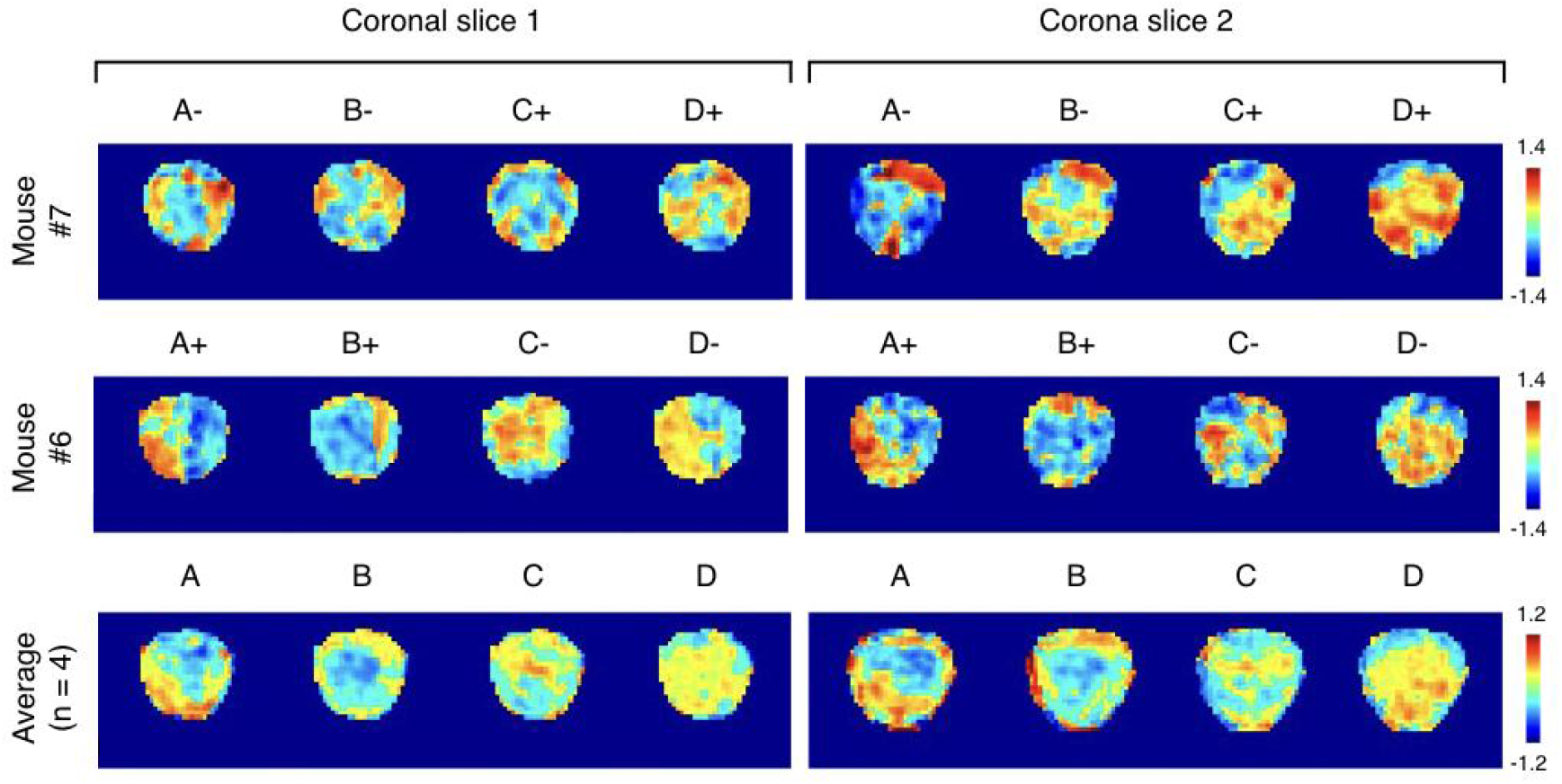
Related to Figure 4 - Odour maps in the olfactory bulb. Averaged maps over the two coronal slices images in the olfactory bulb. **(Top row)** Maps for the example animal shown in the main figure (odours C and D were rewarded). **(Middle row)** Maps for another individual mice with the reverse odour-reward contingencies (odours A and B were rewarded). **(Bottom row)** Average over the 4 mice considered in main Figure 4. Although averaging across mice showed some commonalities, we observed substantial inter-subject variability. This variability is likely explained by a combination of biological and methodological factors. Biologically, some variability is expected due to inter-subject differences in glomeruli and vascular position (ref). Methodologically, we likely introduced substantial variability with (1) differences in slice positioning when imaging different mice and (2) the fact that we counterbalanced reward contingencies across animals, which also modulate the OB (Kay et al., 1999, Doucette et al., 2011). We thus restricted our analyses to within-subject comparisons.

**Figure S4.**
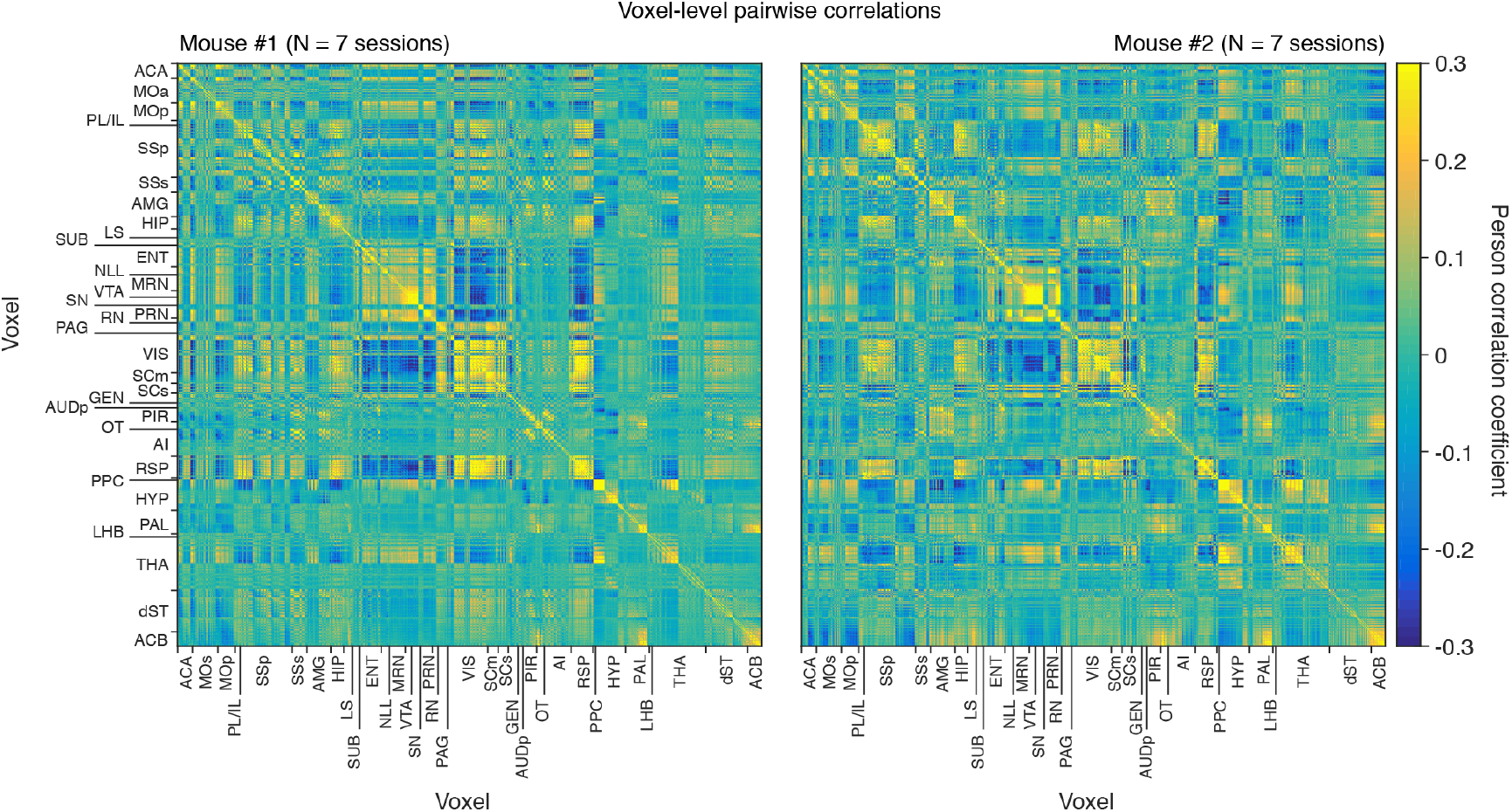
Related to Figure 8 - Voxel-wise functional coupling. Pairwise Pearson correlation coefficients voxel-by-voxel for mouse #1 (left) and mouse #2 (right) in the lever press operant conditioning task. Coefficients were calculated per session (N = 7 sessions), per animal and then averaged across sessions.

**Figure S5.**
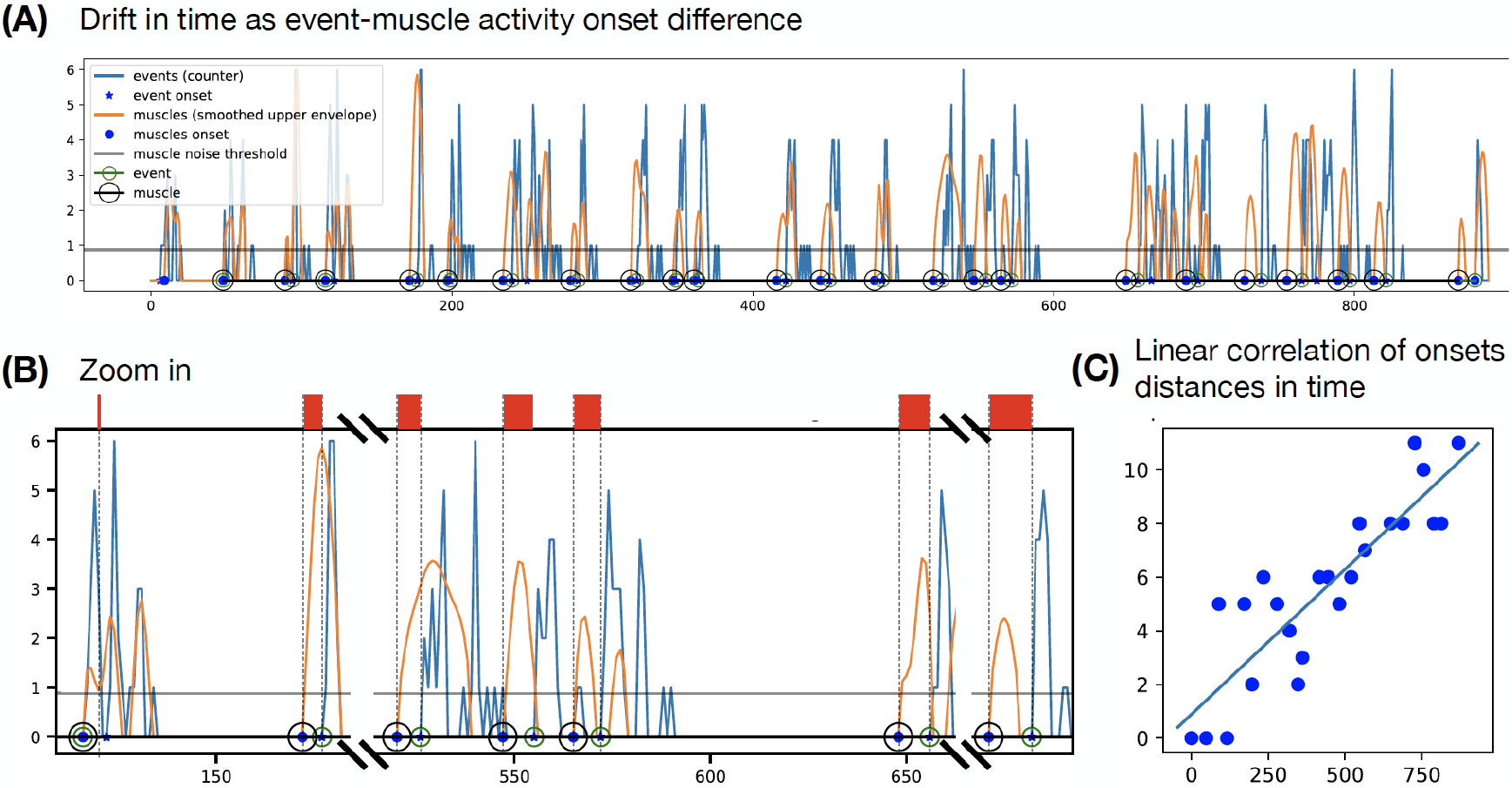
Linear temporal drift correction 1. **(A)** When compared to detected behavioural events (blue time series), the mean of the absolute value of the muscle activity in time (orange) reveals a temporal drift of the Arduino clock with respect to the MR one. **(B)** The delay between the two clocks accumulates in time. **(C)** The delay accumulation in time can be modelled linearly.

**Figure S6.**
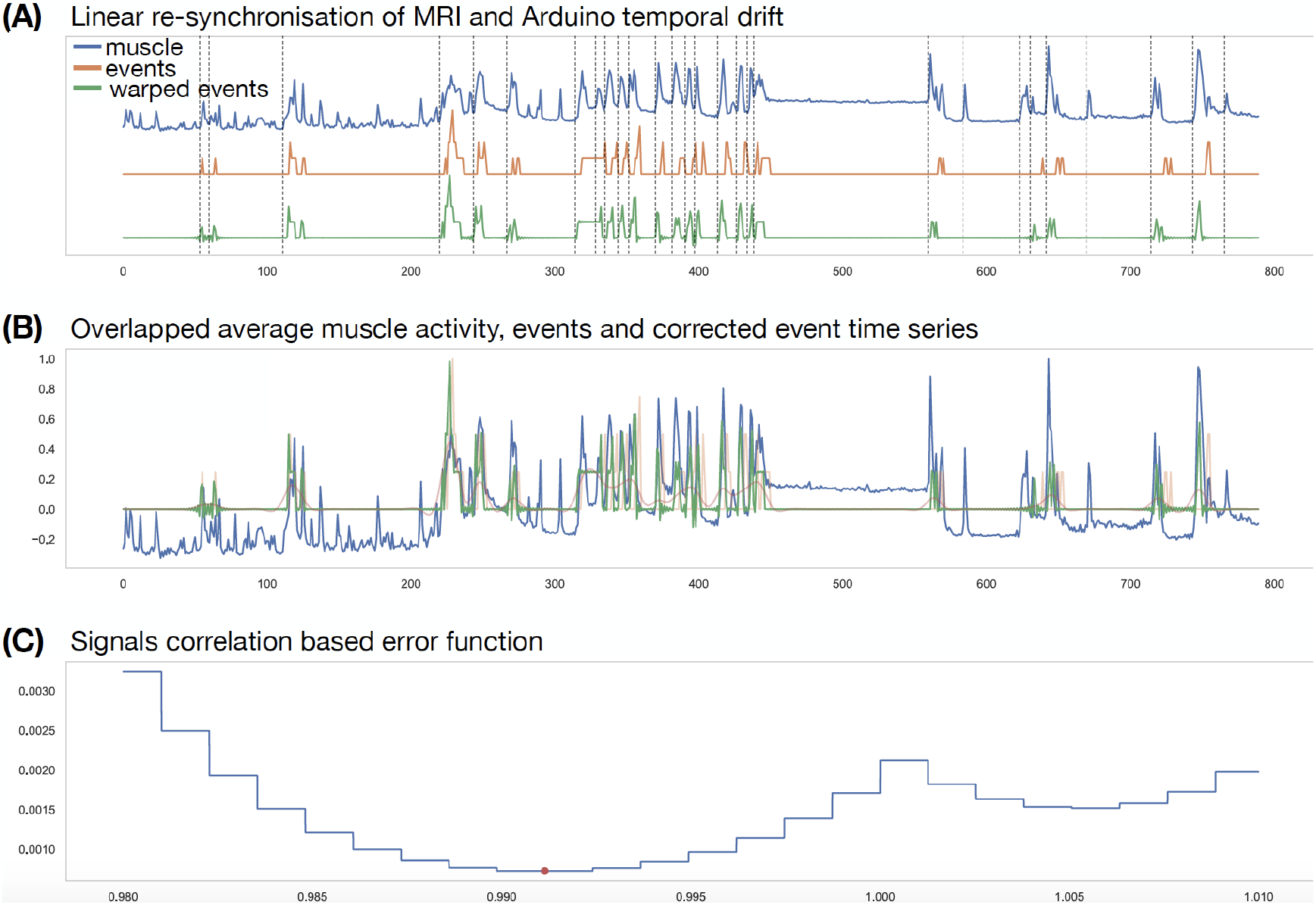
Linear temporal drift correction 2. **(A)** The average of the absolute value of the muscle activity in time (blue time series) is highly correlated with the event detection performed via an Arduino board. The time series of events detected by the Arduino board is depicted as the orange time series. The temporal drift between the two clocks is corrected by maximising the correlation between the two signals and consequently linearly warping the event time series (green). **(B)** as (A). **(C)** The loss function between the muscle time series *M* and the candidate warped event time series *E*(*m̅*) is computed as *L*(*m̅*) = exp (−corr_0_ (*M*, *E* (*m̅*))), where coor_0_ is the operator outputting the correlation at time 0 between the two time series. The value *m* realising the minimum of *L* in the interval [0.98, 1.01] is the warping coefficient applied to correct the temporal drift.

**Figure S7.**
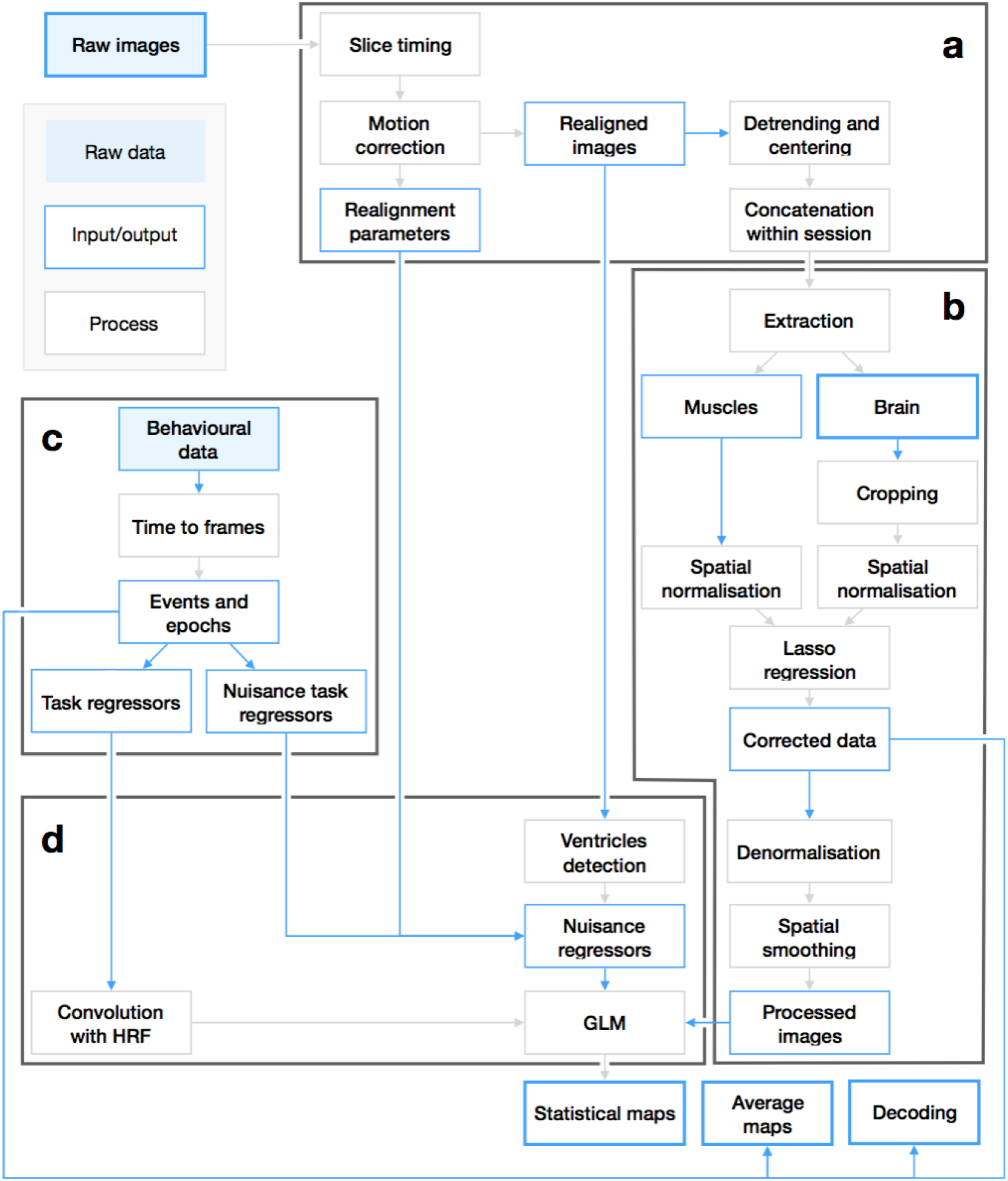
Analysis pipeline: processing of raw images and behavioural data. **(A)** Image preprocessing. Slice timing and motion correction are performed using SPM. Signal is linearly detrended in time and centered with custom Matlab scripts. **(B)** Task-agnostic artefact correction. Masks associated to brain *B_i_* and muscles *M_i_* respectively are manually generated for each slice. Thereafter, images are spatially normalised (2D standardisation). A multivariate LASSO regression is used independently on each slice *s_i_* to compute the contribution of each voxel belonging to *M_i_* to the voxels in *B_i_*. Images are then de-normalised and smoothed spatially with a Gaussian filter. **(C)** Task-related behaviours and epochs are first mapped to the volume acquisition frame rate as boolean vectors or histograms (event occurrences per frame). These vectors are then used as regressor and nuisance regressor in the general linear model used to compute the statistical maps in **(D)**. For every slice, the intensity time course of the brightest pixels corresponding to the ventricles are also used as nuisance regressors.

**Figure S8.**
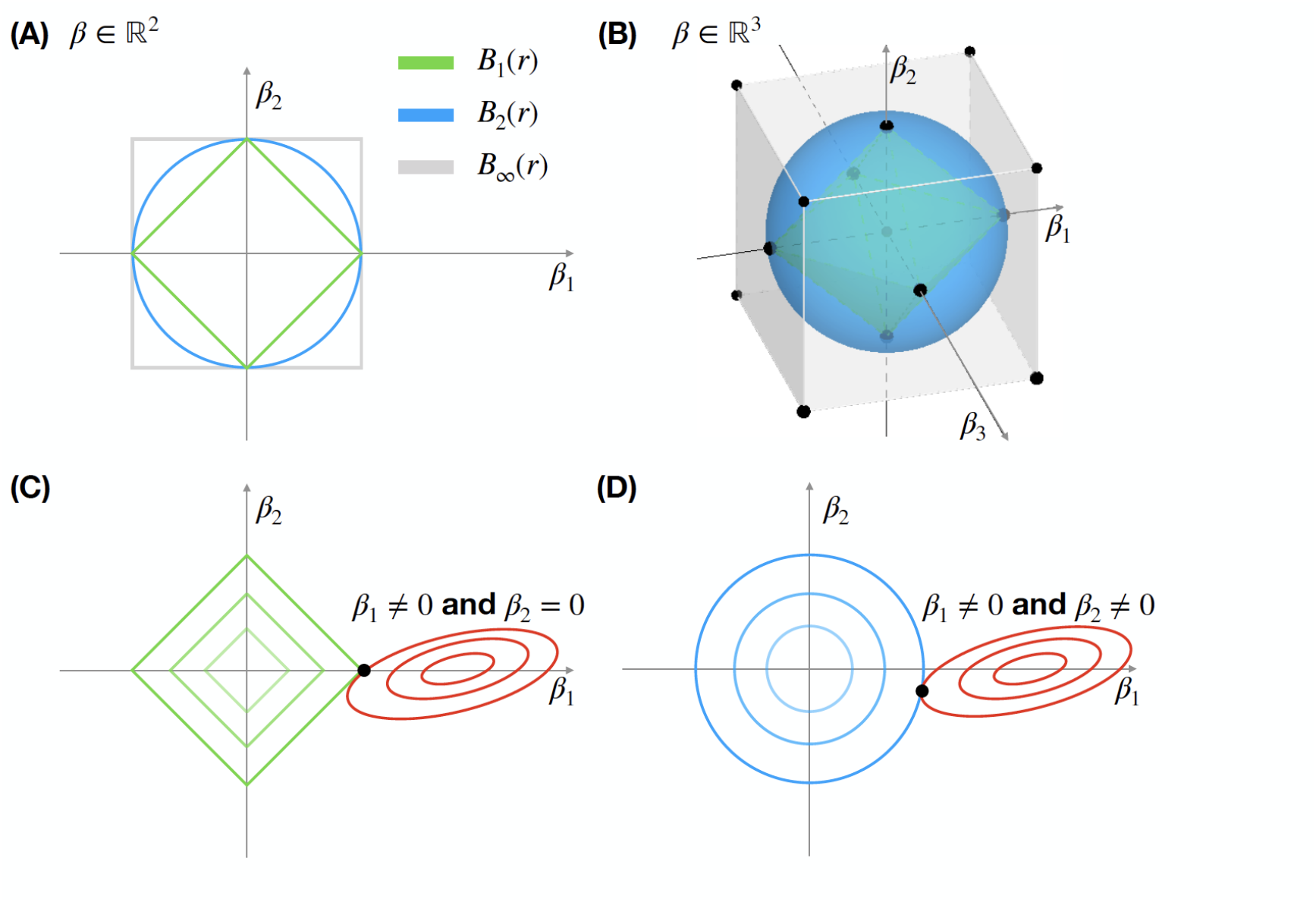
L1 regularisation for voxel selection. **(A)** Different norms induce balls with different shapes. We consider the 1, 2 and ∞− and norms and show the geometries they induce for coefficients in ℛ2 (left) and ℛ_3_ (right), respectively. In symbols we have:

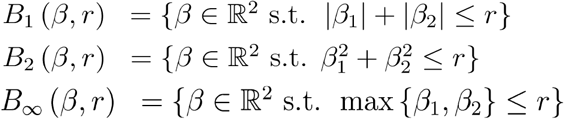

It is important to notice how the ball corresponding to the 1- norm corresponds to a cube whose corners lie exactly on the axes. In **(B)**, the red ellipses represent the contours of the least square error function (loss term in eq. 1). The green and blue shapes correspond to different balls in norms 1 and 2 respectively, as modulated by the parameter of the regularisation term in eq. 1.

**Figure S9.**
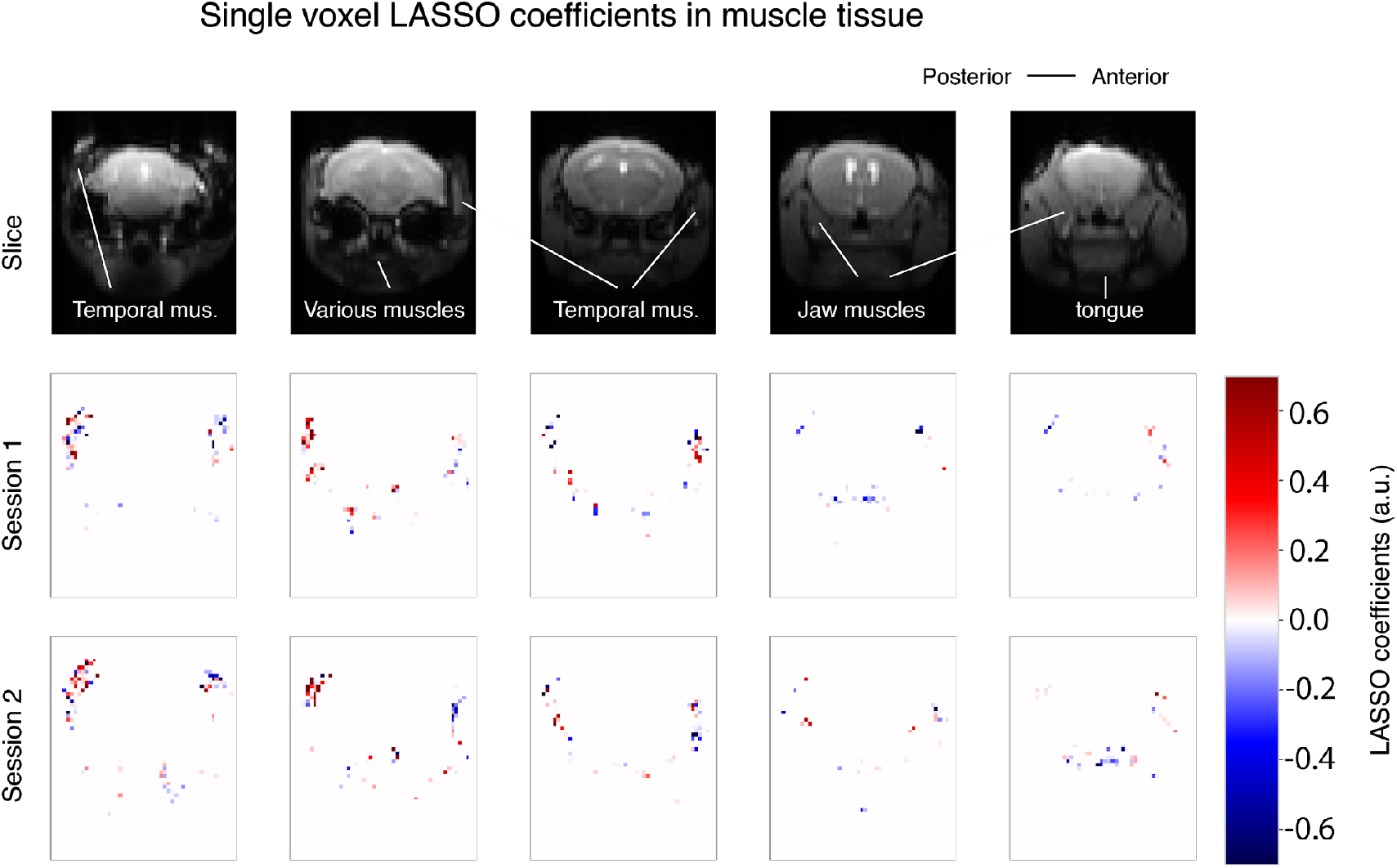
LASSO artefact correction: contribution of each voxel to the predicted intensity of the brain’s voxels. (Top) 5 coronal sections (Middle, bottom) Single voxel LASSO coefficients in the corresponding sections for two example sessions in the lever operant conditioning task (mouse #1). Note that these reflect voxels that were most uniquely predictive, not necessarily all the voxels where muscle movement occurred. This is because LASSO pushes to zero voxels that are redundant or least predictive.

**Figure S10.**
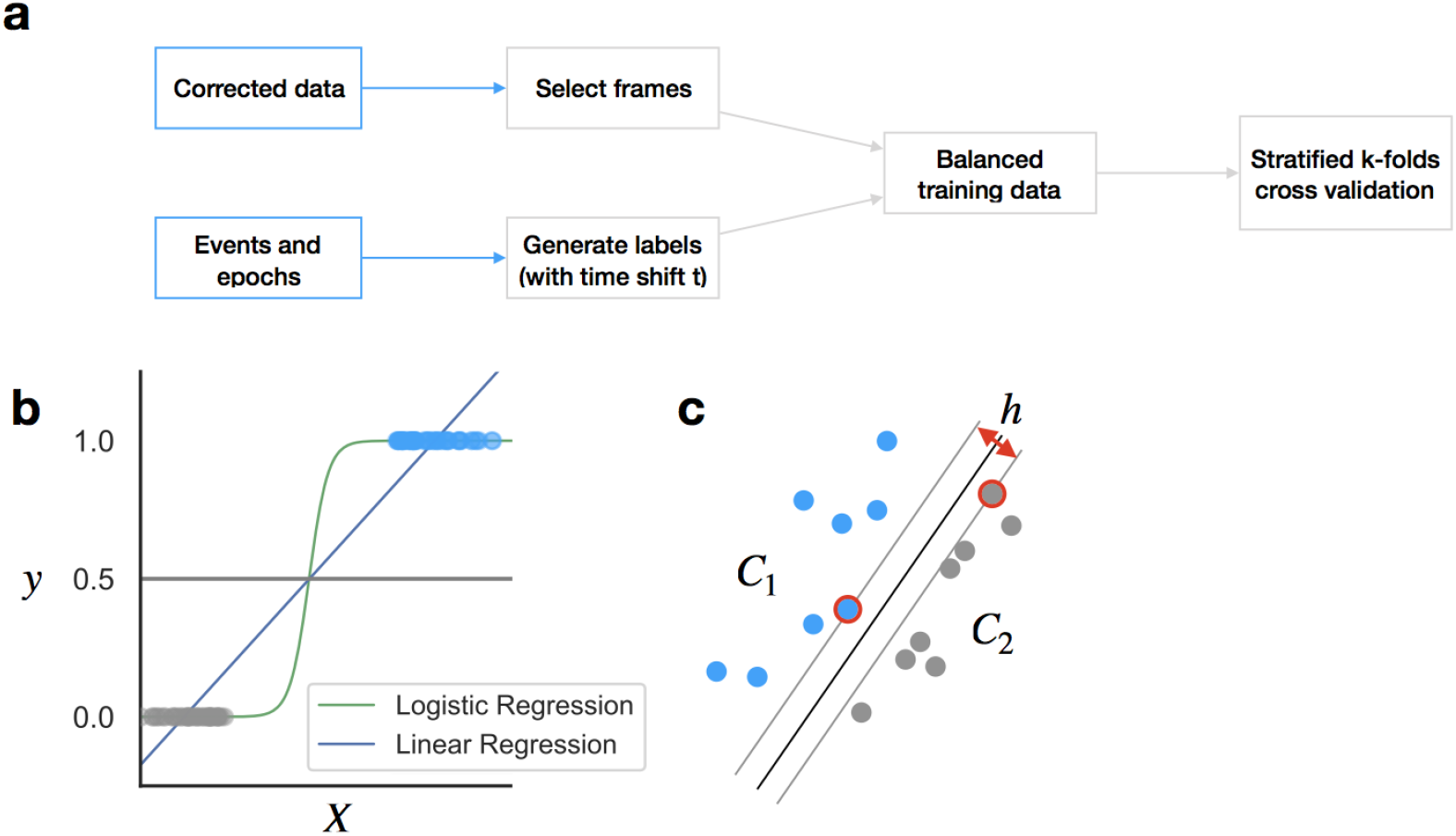
Decoding protocols and algorithms. **(A)** Decoding preprocessing flow diagram. **(B)** Logistic regression. **(C)** Maximal margin hyperplane for separating samples belonging to two classes, as computed by a support vector machine.

**Table S1.** Brain regions mentioned in the text.

| Abbreviation | Full name | Abbreviation | Full name |
| --- | --- | --- | --- |
| ACA | Anterior cingulate area | PH | Posterior hypothalamus |
| ACB | Nucleus Accumbens | PIR | Piriform cortex |
| AI | Agranular insula area | PL | Prelimbic cortex |
| AMG | Amygdala | PO | Preoptic areas (including MPN and MPO) |
| CAMG | Central amygdala | PPC | Posterior parietal cortex |
| dISt | Dorsolateral striatum | PRN | Pontine reticular nucleus |
| dmST | Dorsomedial striatum | PVH | Paraventricular hypothalamic nucleus |
| dST | Dorsal striatum | RN | Red nucleus |
| ENT | Entorhinal cortex | RSP | Retrosplenial cortex |
| GENTHA | Geniculate group of the thalamus | SC | Superior Colliculus |
| HIP | Hippocampus | SCm | Superior Colliculus, motor related |
| HYP | Hypothalamus | SCs | Superior colliculus, sensory related |
| HYP | Hypothalamus | SN | Substantia Nigra |
| IL | Infralimbic cortex | SSp | Primary somatosensory cortex |
| LH | Lateral habenula | SSp(M) | Primary somatosensory cortex, mouth region |
| LHA | Lateral hypothalamic area | SSp(UL) | Primary somatosensory cortex, upper limb region |
| LM | Mamillary nucleus | SSp(UL)L | Primary somatosensory cortex, upper-limb, left |
| LPO | Lateral preoptic area | SSp(UL)R | Primary somatosensory cortex, upper-limb, right |
| LS | Lateral septum | SSs | Supplemental somatosensory cortex |
| MDTHA | Mediodorsal thalamus | SUB | Subiculum |
| MOp | Primary motor cortex | THA | Thalamus |
| MOs | Secondary motor cortex | TRN | Tegmental reticular nucleus |
| MPN | Medial Preoptic nucleus | TT | Taenia Tecta |
| MPO | Medial/median preoptic area | VALTHA | Ventral anterior-lateral complex of the thalamus |
| MRN | Midbrain reticular nucleus | VIS | Visual areas |
| NDB | Diagonal band nucleus | VISp | Primary visual area |
| NLL | Nucleus of the lateral lemniscus | VISpL | Primary visual area, left side |
| OB | Olfactory bulb | VISpR | Primary visual area, right side |
| OT | Olfactory tubercle | VMTHA | Ventromedial thalamus |
| PAG | Periaqueductal gray | VTA | Ventral tegmental area |
| PAL | Pallidum |  |  |

## REFERENCES

Abbott, S. B. G., Machado, N. L. S., Geerling, J. C., & Saper, C. B. (2016). Reciprocal Control of Drinking Behavior by Median Preoptic Neurons in Mice. Journal of Neuroscience, 36(31), 8228–8237. https://doi.org/10.1523/JNEUROSCI.1244-16.2016

Adcock, R. A., Thangavel, A., Whitfield-Gabrieli, S., Knutson, B., & Gabrieli, J. D. E. (2006). Reward-Motivated Learning: Mesolimbic Activation Precedes Memory Formation. Neuron, 50(3), 507–517. https://doi.org/10.1016/j.neuron.2006.03.036

Allen, W. E., DeNardo, L. A., Chen, M. Z., Liu, C. D., Loh, K. M., Fenno, L. E., Ramakrishnan, C., Deisseroth, K., & Luo, L. (2017). Thirst-associated preoptic neurons encode an aversive motivational drive. Science, 357(6356), 1149–1155. https://doi.org/10.1126/science.aan6747

Andersen, R. A., & Cui, H. (2009). Intention, Action Planning, and Decision Making in Parietal-Frontal Circuits. Neuron, 63(5), 568–583. https://doi.org/10.1016/j.neuron.2009.08.028

Blaha, C. D., Yang, C. R., Floresco, S. B., Barr, A. M., & Phillips, A. G. (1997). Stimulation of the Ventral Subiculum of the Hippocampus Evokes Glutamate Receptor-mediated Changes in Dopamine Efflux in the Rat Nucleus Accumbens. European Journal of Neuroscience, 9(5), 902–911. https://doi.org/10.1111/j.1460-9568.1997.tb01441.x

Böhning, D. (1992). Multinomial logistic regression algorithm. Annals of the Institute of Statistical Mathematics, 44(1), 197–200. https://doi.org/10.1007/BF00048682

Boser, B. E., Guyon, I. M., & Vapnik, V. N. (1992). A training algorithm for optimal margin classifiers. Proceedings of the Fifth Annual Workshop on Computational Learning Theory, 144–152. https://doi.org/10.1145/130385.130401

Bradfield, L. A., Hart, G., & Balleine, B. W. (2013). The role of the anterior, mediodorsal, and parafascicular thalamus in instrumental conditioning. Frontiers in Systems Neuroscience, 7. https://doi.org/10.3389/fnsys.2013.00051

Brudzynski, S. M., & Gibson, C. J. (1997). Release of Dopamine in the Nucleus Accumbens Caused By Stimulation of the Subiculum in Freely Moving Rats. Brain Research Bulletin, 42(4), 303–308. https://doi.org/10.1016/S0361-9230(96)00290-0

Burton, H. (1999). Tactile Attention Tasks Enhance Activation in Somatosensory Regions of Parietal Cortex: A Positron Emission Tomography Study. Cerebral Cortex, 9(7), 662–674. https://doi.org/10.1093/cercor/9.7.662

Calu, D. J., Roesch, M. R., Stalnaker, T. A., & Schoenbaum, G. (2007). Associative Encoding in Posterior Piriform Cortex during Odor Discrimination and Reversal Learning. Cerebral Cortex, 17(6), 1342–1349. https://doi.org/10.1093/cercor/bhl045

Cardinal, R. N., Parkinson, J. A., Hall, J., & Everitt, B. J. (2002). Emotion and motivation: The role of the amygdala, ventral striatum, and prefrontal cortex. Neuroscience & Biobehavioral Reviews, 26(3), 321–352. https://doi.org/10.1016/S0149-7634(02)00007-6

Chakraborty, S., Kolling, N., Walton, M. E., & Mitchell, A. S. (2016). Critical role for the mediodorsal thalamus in permitting rapid reward-guided updating in stochastic reward environments. ELife, 5, e13588. https://doi.org/10.7554/eLife.13588

Chapman, C. E., & Meftah, E.-M. (2005). Independent Controls of Attentional Influences in Primary and Secondary Somatosensory Cortex. Journal of Neurophysiology, 94(6), 4094–4107. https://doi.org/10.1152/jn.00303.2005

Cohen, J. Y., Amoroso, M. W., & Uchida, N. (2015). Serotonergic neurons signal reward and punishment on multiple timescales. ELife, 4, e06346. https://doi.org/10.7554/eLife.06346

Cohen, J. Y., Haesler, S., Vong, L., Lowell, B. B., & Uchida, N. (2012). Neuron-type-specific signals for reward and punishment in the ventral tegmental area. Nature, 482(7383), 85–88. https://doi.org/10.1038/nature10754

Cohen, Y. E., & Andersen, R. A. (2002). A common reference frame for movement plans in the posterior parietal cortex. Nature Reviews Neuroscience, 3(7), 553–562. https://doi.org/10.1038/nrn873

Cooper, B. G., & Mizumori, S. J. Y. (2001). Temporary Inactivation of the Retrosplenial Cortex Causes a Transient Reorganization of Spatial Coding in the Hippocampus. The Journal of Neuroscience, 21(11), 3986–4001. https://doi.org/10.1523/JNEUROSCI.21-11-03986.2001

Coq, J.-O., & Xerri, C. (1998). Environmental enrichment alters organizational features of the forepaw representation in the primary somatosensory cortex of adult rats. Experimental Brain Research, 121(2), 191–204. https://doi.org/10.1007/s002210050452

de Quervain, D. J.-F., Roozendaal, B., & McGaugh, J. L. (1998). Stress and glucocorticoids impair retrieval of long-term spatial memory. Nature, 394(6695), 787–790. https://doi.org/10.1038/29542

Debowska, W., Liguz-Lecznar, M., & Kossut, M. (2011). Bilateral Plasticity of Vibrissae SII Representation Induced by Classical Conditioning in Mice. Journal of Neuroscience, 31(14), 5447–5453. https://doi.org/10.1523/JNEUROSCI.5989-10.2011

Deffieux, T., Demene, C., Pernot, M., & Tanter, M. (2018). Functional ultrasound neuroimaging: A review of the preclinical and clinical state of the art. Current Opinion in Neurobiology, 50, 128–135. https://doi.org/10.1016/j.conb.2018.02.001

Desai, M., Kahn, I., Knoblich, U., Bernstein, J., Atallah, H., Yang, A., Kopell, N., Buckner, R. L., Graybiel, A. M., Moore, C. I., & Boyden, E. S. (2011). Mapping brain networks in awake mice using combined optical neural control and fMRI. Journal of Neurophysiology, 105(3), 1393–1405. https://doi.org/10.1152/jn.00828.2010

Desjardins, M., Kılıç, K., Thunemann, M., Mateo, C., Holland, D., Ferri, C. G. L., Cremonesi, J. A., Li, B., Cheng, Q., Weldy, K. L., Saisan, P. A., Kleinfeld, D., Komiyama, T., Liu, T. T., Bussell, R., Wong, E. C., Scadeng, M., Dunn, A. K., Boas, D. A., … Devor, A. (2019). Awake Mouse Imaging: From Two-Photon Microscopy to Blood Oxygen Level–Dependent Functional Magnetic Resonance Imaging. Biological Psychiatry: Cognitive Neuroscience and Neuroimaging, 4(6), 533–542. https://doi.org/10.1016/j.bpsc.2018.12.002

Diamond, D. M., Fleshner, M., & Rose, G. M. (1994). Psychological stress repeatedly blocks hippocampal primed burst potentiation in behaving rats. Behavioural Brain Research, 62(1), 1–9. https://doi.org/10.1016/0166-4328(94)90032-9

Dias-Ferreira, E., Sousa, J. C., Melo, I., Morgado, P., Mesquita, A. R., Cerqueira, J. J., Costa, R. M., & Sousa, N. (2009). Chronic Stress Causes Frontostriatal Reorganization and Affects Decision-Making. Science, 325(5940), 621–625. https://doi.org/10.1126/science.1171203

Doucette, W., Gire, D. H., Whitesell, J., Carmean, V., Lucero, M. T., & Restrepo, D. (2011). Associative Cortex Features in the First Olfactory Brain Relay Station. Neuron, 69(6), 1176–1187. https://doi.org/10.1016/j.neuron.2011.02.024

Eichenbaum, H., Dudchenko, P., Wood, E., Shapiro, M., & Tanila, H. (1999). The Hippocampus, Memory, and Place Cells: Is It Spatial Memory or a Memory Space? Neuron, 23(2), 209–226. https://doi.org/10.1016/S0896-6273(00)80773-4

Elduayen, C., & Save, E. (2014). The retrosplenial cortex is necessary for path integration in the dark. Behavioural Brain Research, 272, 303–307. https://doi.org/10.1016/j.bbr.2014.07.009

Erlich, J. C., Bialek, M., & Brody, C. D. (2011). A Cortical Substrate for Memory-Guided Orienting in the Rat. Neuron, 72(2), 330–343. https://doi.org/10.1016/j.neuron.2011.07.010

Ferenczi, E. A., Zalocusky, K. A., Liston, C., Grosenick, L., Warden, M. R., Amatya, D., Katovich, K., Mehta, H., Patenaude, B., Ramakrishnan, C., Kalanithi, P., Etkin, A., Knutson, B., Glover, G. H., & Deisseroth, K. (2016). Prefrontal cortical regulation of brainwide circuit dynamics and reward-related behavior. Science, 351(6268), aac9698–aac9698. https://doi.org/10.1126/science.aac9698

Friston, K., Ashburner, J., Kiebel, S., Nichols, T., & Penny, W. (2007). Statistical Parametric Mapping. The analysis of Functional Brain Images. Elsevier. https://doi.org/10.1016/B978-0-12-372560-8.X5000-1

Fujiwara, N., Imai, M., Nagamine, T., Mima, T., Oga, T., Takeshita, K., Toma, K., & Shibasaki, H. (2002). Second somatosensory area (SII) plays a significant role in selective somatosensory attention. Cognitive Brain Research, 14(3), 389–397. https://doi.org/10.1016/S0926-6410(02)00141-6

Glover, G. H. (1999). Deconvolution of Impulse Response in Event-Related BOLD fMRI1. NeuroImage, 9(4), 416–429. https://doi.org/10.1006/nimg.1998.0419

Good, M. (2002). Spatial Memory and Hippocampal Function: Where are we now? Psicologica, 23, 109–138.

Graham, L. K., Yoon, T., & Kim, J. J. (2010). Stress impairs optimal behavior in a water foraging choice task in rats. Learning & Memory, 17(1), 1–4. https://doi.org/10.1101/lm.1605510

Guenthner, C. J., Miyamichi, K., Yang, H. H., Heller, H. C., & Luo, L. (2013). Permanent Genetic Access to Transiently Active Neurons via TRAP: Targeted Recombination in Active Populations. Neuron, 78(5), 773–784. https://doi.org/10.1016/j.neuron.2013.03.025

Guo, Z. V., Hires, S. A., Li, N., O’Connor, D. H., Komiyama, T., Ophir, E., Huber, D., Bonardi, C., Morandell, K., Gutnisky, D., Peron, S., Xu, N., Cox, J., & Svoboda, K. (2014). Procedures for Behavioral Experiments in Head-Fixed Mice. PLoS ONE, 9(2), e88678. https://doi.org/10.1371/journal.pone.0088678

Guo, Z. V., Li, N., Huber, D., Ophir, E., Gutnisky, D., Ting, J. T., Feng, G., & Svoboda, K. (2014). Flow of Cortical Activity Underlying a Tactile Decision in Mice. Neuron, 81(1), 179–194. https://doi.org/10.1016/j.neuron.2013.10.020

Haberly, L. B. (2001). Parallel-distributed Processing in Olfactory Cortex: New Insights from Morphological and Physiological Analysis of Neuronal Circuitry. Chemical Senses, 26(5), 551–576. https://doi.org/10.1093/chemse/26.5.551

Han, Z., Chen, W., Chen, X., Zhang, K., Tong, C., Zhang, X., Li, C. T., & Liang, Z. (2019). Awake and behaving mouse fMRI during Go/No-Go task. NeuroImage, 188, 733–742. https://doi.org/10.1016/j.neuroimage.2019.01.002

Harris, A. P., Lennen, R. J., Marshall, I., Jansen, M. A., Pernet, C. R., Brydges, N. M., Duguid, I. C., & Holmes, M. C. (2015). Imaging learned fear circuitry in awake mice using fMRI. European Journal of Neuroscience, 42(5), 2125–2134. https://doi.org/10.1111/ejn.12939

Hölscher, C. (1999). Stress impairs performance in spatial water maze learning tasks. Behavioural Brain Research, 100(1–2), 225–235. https://doi.org/10.1016/S0166-4328(98)00134-X

Hoover, W. B., & Vertes, R. P. (2011). Projections of the medial orbital and ventral orbital cortex in the rat. The Journal of Comparative Neurology, 519(18), 3766–3801. https://doi.org/10.1002/cne.22733

Huttunen, J., Wikström, H., Korvenoja, A., Seppäläinen, A.-M., Aronen, H., & llmoniemi, R. J. (1996). Significance of the second somatosensory cortex in sensorimotor integration: Enhancement of sensory responses during finger movements. Neuroreport: An International Journal for the Rapid Communication of Research in Neuroscience, 7(5), 1009–1012. https://doi.org/10.1097/00001756-199604100-00011

Igarashi, K. M., Ieki, N., An, M., Yamaguchi, Y., Nagayama, S., Kobayakawa, K., Kobayakawa, R., Tanifuji, M., Sakano, H., Chen, W. R., & Mori, K. (2012). Parallel Mitral and Tufted Cell Pathways Route Distinct Odor Information to Different Targets in the Olfactory Cortex. Journal of Neuroscience, 32(23), 7970–7985. https://doi.org/10.1523/JNEUROSCI.0154-12.2012

Izquierdo, A., & Murray, E. A. (2010). Functional Interaction of Medial Mediodorsal Thalamic Nucleus But Not Nucleus Accumbens with Amygdala and Orbital Prefrontal Cortex Is Essential for Adaptive Response Selection after Reinforcer Devaluation. Journal of Neuroscience, 30(2), 661–669. https://doi.org/10.1523/JNEUROSCI.3795-09.2010

Janak, P. H., & Tye, K. M. (2015). From circuits to behaviour in the amygdala. Nature, 517(7534), 284–292. https://doi.org/10.1038/nature14188

Ji, L. L., Fleming, T., Penny, M. L., Toney, G. M., & Cunningham, J. T. (2005). Effects of water deprivation and rehydration on c-Fos and FosB staining in the rat supraoptic nucleus and lamina terminalis region. American Journal of Physiology-Regulatory, Integrative and Comparative Physiology, 288(1), R311–R321. https://doi.org/10.1152/ajpregu.00399.2004

Kaneto, H. (1997). Learning/memory processes under stress conditions. Behavioural Brain Research, 83(1–2), 71–74. https://doi.org/10.1016/S0166-4328(97)86048-2

Kawai, R., Markman, T., Poddar, R., Ko, R., Fantana, A. L., Dhawale, A. K., Kampff, A. R., & Ölveczky, B. P. (2015). Motor Cortex Is Required for Learning but Not for Executing a Motor Skill. Neuron, 86(3), 800–812. https://doi.org/10.1016/j.neuron.2015.03.024

Kay, L. M., & Laurent, G. (1999). Odor- and context-dependent modulation of mitral cell activity in behaving rats. Nature Neuroscience, 2(11), 1003–1009. https://doi.org/10.1038/14801

Keliris, G. A., Shmuel, A., Ku, S.-P., Pfeuffer, J., Oeltermann, A., Steudel, T., & Logothetis, N. K. (2007). Robust controlled functional MRI in alert monkeys at high magnetic field: Effects of jaw and body movements. NeuroImage, 36(3), 550–570. https://doi.org/10.1016/j.neuroimage.2007.02.057

Kepecs, A., Uchida, N., & Mainen, Z. F. (2007). Rapid and Precise Control of Sniffing During Olfactory Discrimination in Rats. Journal of Neurophysiology, 98(1), 205–213. https://doi.org/10.1152/jn.00071.2007

Kim, J. J., Lee, H. J., Han, J.-S., & Packard, M. G. (2001). Amygdala Is Critical for Stress-Induced Modulation of Hippocampal Long-Term Potentiation and Learning. The Journal of Neuroscience, 21(14), 5222–5228. https://doi.org/10.1523/JNEUROSCI.21-14-05222.2001

King, J. A., Garelick, T. S., Brevard, M. E., Chen, W., Messenger, T. L., Duong, T. Q., & Ferris, C. F. (2005). Procedure for minimizing stress for fMRI studies in conscious rats. Journal of Neuroscience Methods, 148(2), 154–160. https://doi.org/10.1016/j.jneumeth.2005.04.011

Kolling, N., Wittmann, M. K., Behrens, T. E. J., Boorman, E. D., Mars, R. B., & Rushworth, M. F. S. (2016). Value, search, persistence and model updating in anterior cingulate cortex. Nature Neuroscience, 19(10), 1280–1285. https://doi.org/10.1038/nn.4382

Kononenko, N. L., & Witter, M. P. (2012). Presubiculum layer III conveys retrosplenial input to the medial entorhinal cortex. Hippocampus, 22(4), 881–895. https://doi.org/10.1002/hipo.20949

Korsching, S. I. (2001). Odor maps in the brain: Spatial aspects of odor representation in sensory surface and olfactory bulb: Cellular and Molecular Life Sciences, 58(4), 520–530. https://doi.org/10.1007/PL00000877

Krakauer, J. W., Hadjiosif, A. M., Xu, J., Wong, A. L., & Haith, A. M. (2019). Motor Learning. Comprehensive Physiology, 9(2), 613–663. https://doi.org/10.1002/cphy.c170043.

Laing, D. G. (1983). Natural Sniffing Gives Optimum Odour Perception for Humans. Perception, 12(2), 99–117. https://doi.org/10.1068/p120099

Legault, M., Rompré, P.-P., & Wise, R. A. (2000). Chemical Stimulation of the Ventral Hippocampus Elevates Nucleus Accumbens Dopamine by Activating Dopaminergic Neurons of the Ventral Tegmental Area. The Journal of Neuroscience, 20(4), 1635–1642. https://doi.org/10.1523/JNEUROSCI.20-04-01635.2000

Legault, M., & Wise, R. A. (1999). Injections of N-methyl-D-aspartate into the ventral hippocampus increase extracellular dopamine in the ventral tegmental area and nucleus accumbens. Synapse, 31, 241–249.

Leon, M., & Johnson, B. A. (2003). Olfactory coding in the mammalian olfactory bulb. Brain Research Reviews, 42(1), 23–32. https://doi.org/10.1016/S0165-0173(03)00142-5

Li, N., Chen, T.-W., Guo, Z. V., Gerfen, C. R., & Svoboda, K. (2015). A motor cortex circuit for motor planning and movement. Nature, 519(7541), 51–56. https://doi.org/10.1038/nature14178

Lisman, J. E., & Grace, A. A. (2005). The Hippocampal-VTA Loop: Controlling the Entry of Information into Long-Term Memory. Neuron, 46(5), 703–713. https://doi.org/10.1016/j.neuron.2005.05.002

Lu, R., Liang, Y., Meng, G., Zhou, P., Svoboda, K., Paninski, L., & Ji, N. (2020). Rapid mesoscale volumetric imaging of neural activity with synaptic resolution. Nature Methods, 17(3), 291–294. https://doi.org/10.1038/s41592-020-0760-9

Luft, A. R., & Buitrago, M. M. (2005). Stages of Motor Skill Learning. Molecular Neurobiology, 32(3), 205–216. https://doi.org/10.1385/MN:32:3:205

Macé, É., Montaldo, G., Trenholm, S., Cowan, C., Brignall, A., Urban, A., & Roska, B. (2018). Whole-Brain Functional Ultrasound Imaging Reveals Brain Modules for Visuomotor Integration. Neuron, 100(5), 1241–1251.e7. https://doi.org/10.1016/j.neuron.2018.11.031

Mair, R. G., Miller, R. L. A., Wormwood, B. A., Francoeur, M. J., Onos, K. D., & Gibson., B. M. (2015). The neurobiology of thalamic amnesia: Contributions of medial thalamus and prefrontal cortex to delayed conditional discrimination. Neuroscience & Biobehavioral Reviews, 54, 161–174. https://doi.org/10.1016/j.neubiorev.2015.01.011

Manzoni, T., Caminiti, R., Spidalieri, G., & Morelli, E. (1979). Anatomical and functional aspects of the associative projections from somatic area SI to SII. Experimental Brain Research, 34(3). https://doi.org/10.1007/BF00239142

Martin, P. (2001). Locomotion towards a goal alters the synchronous firing of neurons recorded simultaneously in the subiculum and nucleus accumbens of rats. Behavioural Brain Research, 124(1), 19–28. https://doi.org/10.1016/S0166-4328(01)00209-1

Marton, T. F., Seifikar, H., Luongo, F. J., Lee, A. T., & Sohal, V. S. (2018). Roles of Prefrontal Cortex and Mediodorsal Thalamus in Task Engagement and Behavioral Flexibility. The Journal of Neuroscience, 38(10), 2569–2578. https://doi.org/10.1523/JNEUROSCI.1728-17.2018

Matias, S., Lottem, E., Dugué, G. P., & Mainen, Z. F. (2017). Activity patterns of serotonin neurons underlying cognitive flexibility. ELife, 6, e20552. https://doi.org/10.7554/eLife.20552

McKinley, M. J., Hards, D. K., & Oldfield, B. J. (1994). Identification of neural pathways activated in dehydrated rats by means of Fos-immunohistochemistry and neural tracing. Brain Research, 653(1–2), 305–314. https://doi.org/10.1016/0006-8993(94)90405-7

McNaughton, B. L., & Morris, R. G. M. (1987). Hippocampal synaptic enhancement and information storage within a distributed memory system. Trends in Neurosciences, 10(10), 8.

Meftah, E.-M., Shenasa, J., & Chapman, C. E. (2002). Effects of a Cross-Modal Manipulation of Attention on Somatosensory Cortical Neuronal Responses to Tactile Stimuli in the Monkey. Journal of Neurophysiology, 88(6), 3133–3149. https://doi.org/10.1152/jn.00121.2002

Mesulam, M.-M., Mufson, E. J., Wainer, B. H., & Levey, A. I. (1983). Central cholinergic pathways in the rat: An overview based on an alternative nomenclature (Ch1–Ch6). Neuroscience, 10(4), 1185–1201. https://doi.org/10.1016/0306-4522(83)90108-2

Miller, K. L. (2012). FMRI using balanced steady-state free precession (SSFP). NeuroImage, 62(2), 713–719. https://doi.org/10.1016/j.neuroimage.2011.10.040

Mima, T., Nagamine, T., Nakamura, K., & Shibasaki, H. (1998). Attention Modulates Both Primary and Second Somatosensory Cortical Activities in Humans: A Magnetoencephalographic Study. Journal of Neurophysiology, 80(4), 2215–2221. https://doi.org/10.1152/jn.1998.80.4.2215

Mitchell, A. S. (2015). The mediodorsal thalamus as a higher order thalamic relay nucleus important for learning and decision-making. Neuroscience & Biobehavioral Reviews, 54, 76–88. https://doi.org/10.1016/j.neubiorev.2015.03.001

Mitchell, A. S., & Chakraborty, S. (2013). What does the mediodorsal thalamus do? Frontiers in Systems Neuroscience, 7. https://doi.org/10.3389/fnsys.2013.00037

Miyashita, T., & Rockland, K. S. (2007). GABAergic projections from the hippocampus to the retrosplenial cortex in the rat: Hippocampo-retrosplenial GABAergic projections. European Journal of Neuroscience, 26(5), 1193–1204. https://doi.org/10.1111/j.1460-9568.2007.05745.x

Mori, K. (2003). Grouping of odorant receptors: Odour maps in the mammalian olfactory bulb. Biochemical Society Transactions, 31(1), 134–136. https://doi.org/10.1042/bst0310134

Morris, R. G. M., & Frey, U. (1997). Hippocampal synaptic plasticity: Role in spatial learning or the automatic recording of attended experience? Philosophical Transactions of the Royal Society of London. Series B: Biological Sciences, 352(1360), 1489–1503. https://doi.org/10.1098/rstb.1997.0136

Motta, S. C., Carobrez, A. P., & Canteras, N. S. (2017). The periaqueductal gray and primal emotional processing critical to influence complex defensive responses, fear learning and reward seeking. Neuroscience & Biobehavioral Reviews, 76, 39–47. https://doi.org/10.1016/j.neubiorev.2016.10.012

Murty, V. P., & Adcock, R. A. (2014). Enriched Encoding: Reward Motivation Organizes Cortical Networks for Hippocampal Detection of Unexpected Events. Cerebral Cortex, 24(8), 2160–2168. https://doi.org/10.1093/cercor/bht063

Musall, S., Kaufman, M. T., Juavinett, A. L., Gluf, S., & Churchland, A. K. (2019). Single-trial neural dynamics are dominated by richly varied movements. Nature Neuroscience, 22(10), 1677–1686. https://doi.org/10.1038/s41593-019-0502-4

Ostlund, S. B., & Balleine, B. W. (2008). Differential Involvement of the Basolateral Amygdala and Mediodorsal Thalamus in Instrumental Action Selection. Journal of Neuroscience, 28(17), 4398–4405. https://doi.org/10.1523/JNEUROSCI.5472-07.2008

Pachitariu, M., Stringer, C., & Harris, K. D. (2018). Robustness of Spike Deconvolution for Neuronal Calcium Imaging. The Journal of Neuroscience, 38(37), 7976–7985. https://doi.org/10.1523/JNEUROSCI.3339-17.2018

Park, S.-H., Kim, T., Wang, P., & Kim, S.-G. (2011). Sensitivity and specificity of high-resolution balanced steady-state free precession fMRI at high field of 9.4T. NeuroImage, 58(1), 168–176. https://doi.org/10.1016/j.neuroimage.2011.06.010

Quallo, M. M., Price, C. J., Ueno, K., Asamizuya, T., Cheng, K., Lemon, R. N., & Iriki, A. (2009). Gray and white matter changes associated with tool-use learning in macaque monkeys. Proceedings of the National Academy of Sciences, 106(43), 18379–18384. https://doi.org/10.1073/pnas.0909751106

Rabut, C., Correia, M., Finel, V., Pezet, S., Pernot, M., Deffieux, T., & Tanter, M. (2019). 4D functional ultrasound imaging of whole-brain activity in rodents. Nature Methods, 16(10), 994–997. https://doi.org/10.1038/s41592-019-0572-y

Roesch, M. R., Stalnaker, T. A., & Schoenbaum, G. (2006). Associative Encoding in Anterior Piriform Cortex versus Orbitofrontal Cortex during Odor Discrimination and Reversal Learning. Cerebral Cortex, 17(3), 643–652. https://doi.org/10.1093/cercor/bhk009

Royet, J. P., Souchier, C., Jourdan, F., & Ploye, H. (1988). Morphometric study of the glomerular population in the mouse olfactory bulb: Numerical density and size distribution along the rostrocaudal axis. The Journal of Comparative Neurology, 270(4), 559–568. https://doi.org/10.1002/cne.902700409

Rubin, B. D., & Katz, L. C. (1999). Optical Imaging of Odorant Representations in the Mammalian Olfactory Bulb. Neuron, 23(3), 499–511. https://doi.org/10.1016/S0896-6273(00)80803-X

Sakurai, K., Shintani, T., Jomura, N., Matsuda, T., Sumiyoshi, A., & Hisatsune, T. (2020). Hyper BOLD Activation in Dorsal Raphe Nucleus of APP/PS1 Alzheimer’s Disease Mouse during Reward-Oriented Drinking Test under Thirsty Conditions. Scientific Reports, 10(1), 3915. https://doi.org/10.1038/s41598-020-60894-7

Shohamy, D., & Adcock, R. A. (2010). Dopamine and adaptive memory. Trends in Cognitive Sciences, 14(10), 464–472. https://doi.org/10.1016/j.tics.2010.08.002

Sieu, L.-A., Bergel, A., Tiran, E., Deffieux, T., Pernot, M., Gennisson, J.-L., Tanter, M., & Cohen, I. (2015). EEG and functional ultrasound imaging in mobile rats. Nature Methods, 12(9), 831–834. https://doi.org/10.1038/nmeth.3506

Sobel, N. (2000). Sniffing Longer rather than Stronger to Maintain Olfactory Detection Threshold. Chemical Senses, 25(1), 1–8. https://doi.org/10.1093/chemse/25.1.1

Sofroniew, N. J., Flickinger, D., King, J., & Svoboda, K. (2016). A large field of view two-photon mesoscope with subcellular resolution for in vivo imaging. ELife, 5, e14472. https://doi.org/10.7554/eLife.14472

Steinmetz, N. A., Zatka-Haas, P., Carandini, M., & Harris, K. D. (2019). Distributed coding of choice, action and engagement across the mouse brain. Nature, 576(7786), 266–273. https://doi.org/10.1038/s41586-019-1787-x

Sugar, J., Witter, M. P., van Strien, N. M., & Cappaert, N. L. M. (2011). The Retrosplenial Cortex: Intrinsic Connectivity and Connections with the (Para)Hippocampal Region in the Rat. An Interactive Connectome. Frontiers in Neuroinformatics, 5. https://doi.org/10.3389/fninf.2011.00007

Sul, J. H., Jo, S., Lee, D., & Jung, M. W. (2011). Role of rodent secondary motor cortex in value-based action selection. Nature Neuroscience, 14(9), 1202–1208. https://doi.org/10.1038/nn.2881

Suykens, J. A. K., & Vandewalle, J. (1999). Least Squares Support Vector Machine Classifiers. Neural Processing Letters, 9, 293–300.

Tabuchi, E., Yokawa, T., Mallick, H., Inubushi, T., Kondoh, T., Ono, T., & Torii, K. (2002). Spatio–temporal dynamics of brain activated regions during drinking behavior in rats. Brain Research, 951(2), 270–279. https://doi.org/10.1016/S0006-8993(02)03173-6

Theis, L., Berens, P., Froudarakis, E., Reimer, J., Román Rosón, M., Baden, T., Euler, T., Tolias, A. S., & Bethge, M. (2016). Benchmarking Spike Rate Inference in Population Calcium Imaging. Neuron, 90(3), 471–482. https://doi.org/10.1016/j.neuron.2016.04.014

Tian, J., & Uchida, N. (2015). Habenula Lesions Reveal that Multiple Mechanisms Underlie Dopamine Prediction Errors. Neuron, 87(6), 1304–1316. https://doi.org/10.1016/j.neuron.2015.08.028

Tibshirani, R. (1996). Regression Shrinkage and Selection via the Lasso. Journal of the Royal Statistical Society. Series B (Methodological), 58(1), 267–288. JSTOR.

Tryon, V. L., & Mizumori, S. J. Y. (2018). A Novel Role for the Periaqueductal Gray in Consummatory Behavior. Frontiers in Behavioral Neuroscience, 12, 178. https://doi.org/10.3389/fnbeh.2018.00178

Urban, A., Mace, E., Brunner, C., Heidmann, M., Rossier, J., & Montaldo, G. (2014). Chronic assessment of cerebral hemodynamics during rat forepaw electrical stimulation using functional ultrasound imaging. NeuroImage, 101, 138–149. https://doi.org/10.1016/j.neuroimage.2014.06.063

Van de Moortele, P.-F., Pfeuffer, J., Glover, G. H., Ugurbil, K., & Hu, X. (2002). Respiration-inducedB0 fluctuations and their spatial distribution in the human brain at 7 Tesla. Magnetic Resonance in Medicine, 47(5), 888–895. https://doi.org/10.1002/mrm.10145

van Groen, T., & Wyss, J. M. (1992). Connections of the retrosplenial dysgranular cortex in the rat. The Journal of Comparative Neurology, 315(2), 200–216. https://doi.org/10.1002/cne.903150207

Vann, S. D., & Aggleton, J. P. (2004). The mammillary bodies: Two memory systems in one? Nature Reviews Neuroscience, 5(1), 35–44. https://doi.org/10.1038/nrn1299

Vann, S. D., Aggleton, J. P., & Maguire, E. A. (2009). What does the retrosplenial cortex do? Nature Reviews Neuroscience, 10(11), 792–802. https://doi.org/10.1038/nrn2733

Vassar, R., Chao, S. K., & Sitcheran, R. (1994). Topographic Organization of Sensory Projection to the Olfactory Bulb. Cell, 79, 981–991.

Vogelstein, J. T., Packer, A. M., Machado, T. A., Sippy, T., Babadi, B., Yuste, R., & Paninski, L. (2010). Fast Nonnegative Deconvolution for Spike Train Inference From Population Calcium Imaging. Journal of Neurophysiology, 104(6), 3691–3704. https://doi.org/10.1152/jn.01073.2009

Vorel, S. R. (2001). Relapse to Cocaine-Seeking After Hippocampal Theta Burst Stimulation. Science, 292(5519), 1175–1178. https://doi.org/10.1126/science.1058043

Warren, D. W., Walker, J. C., Drake, A. F., & Lutz, R. W. (1994). Effects of odorants and irritants on respiratory behavior. The Laryngoscope, 104(5), 623–626. https://doi.org/10.1002/lary.5541040517

Wekselblatt, J. B., Flister, E. D., Piscopo, D. M., & Niell, C. M. (2016). Large-scale imaging of cortical dynamics during sensory perception and behavior. Journal of Neurophysiology, 115(6), 2852–2866. https://doi.org/10.1152/jn.01056.2015

Wittmann, B. C., Schott, B. H., Guderian, S., Frey, J. U., Heinze, H.-J., & Düzel, E. (2005). Reward-Related fMRI Activation of Dopaminergic Midbrain Is Associated with Enhanced Hippocampus- Dependent Long-Term Memory Formation. Neuron, 45(3), 459–467. https://doi.org/10.1016/j.neuron.2005.01.010

Youngentob, S. L., Mozell, M. M., Sheehe, P. R., & Hornung, D. E. (1987). A quantitative analysis of sniffing strategies in rats performing odor detection tasks. Physiology & Behavior, 41(1), 59–69. https://doi.org/10.1016/0031-9384(87)90131-4

Zhang, Z., Seginer, A., & Frydman, L. (2017). Single-scan MRI with exceptional resilience to field heterogeneities: Single-Scan Distortion-Free xSPEN MRI. Magnetic Resonance in Medicine, 77(2), 623–634. https://doi.org/10.1002/mrm.26145

Zhou, I. Y., Cheung, M. M., Lau, C., Chan, K. C., & Wu, E. X. (2012). Balanced steady-state free precession fMRI with intravascular susceptibility contrast agent. Magnetic Resonance in Medicine, 68(1), 65–73. https://doi.org/10.1002/mrm.23202

